# Sequential IL-15 and TGF-β1 stimulation expands reprogrammed CD103⁺CD39⁺ effector CD8⁺T cells and enhances HIV control

**DOI:** 10.1101/2025.04.09.647906

**Authors:** Cristina Mancebo-Pérez, Aleix Benítez-Martínez, Ilya Tsukalov, Judith Grau-Expósito, Josep Castellví, Laura Mañalich-Barrachina, Cristina Centeno-Mediavilla, Bibiana Planas, Jordi Navarro, Joaquín Burgos, Paula Suanzes, Adrian Curran, Vicenç Falcó, Enrique Martín-Gayo, María J. Buzón, Meritxell Genescà

## Abstract

During analytical treatment interruption (ATI), failure to contain tissue HIV reservoirs remains a major barrier to remission. We conditioned peripheral blood mononuclear cells with cues that promote tissue-resident memory differentiation, aiming to reprogram HIV- specific CD8^+^T cells for these tissue microenvironments. Sequential IL-15/TGF-β1- stimulation expanded proliferative CD103^+^CD39^+^ effector populations and restored polyfunctional HIV-specific CD8^+^T cell responses, irrespective of CD39 or PD-1 expression. Cytokine-stimulated CD8^+^T cells enhanced *ex vivo* elimination of the HIV reservoir, and within the induced effector pool, only CD69^+^ subsets expressing CD103^+^ or CD39^+^ eliminated intact HIV-1 genomes. Single-cell transcriptomics showed that IL- 15/TGF-β1 stimulation increased clonotypic diversity and expanded resident-like effector clusters with reduced immune-checkpoint receptor expression and augmented mitochondrial function, findings confirmed by flow cytometry. Engrafted recipients from a humanized mouse ATI model more effectively eliminated HIV^+^-reactivated cells after adoptive transfer of sequentially-treated CD8^+^T cells. Thus, IL-15/TGF-β1-mediated CD8^+^T cell reprogramming represents a promising immunotherapy to enable immune control after ATI.

---

The establishment of viral reservoirs remains the major obstacle to achieve a cure for antiretroviral therapy (ART)-treated individuals with human immunodeficiency virus (HIV)^1^. Studies focused on understanding the source of viruses that emerge during ART interruption demonstrate that they are established in different cellular and anatomical compartments, denoting substantial inter-participant variability ^2,3^. Thus, a major challenge lies in the fact that nearly all tissues assessed in people with HIV (PWH) exhibit some level of viral persistence^4^. Moreover, simian immunodeficiency virus (SIV) models have shown that potent antiviral CD8^+^T cell responses occur too late to limit local viral reactivation and prevent the early spread of viral reservoirs upon ART withdrawal^5^. Altogether, these findings suggest the need to promote effective CD8^+^T cells that localize to multiple tissue compartments to control reactivated viruses; perhaps as CD8^+^ tissue resident memory T cell (TRM) phenotypes. The CD8^+^TRM pool reside within and actively survey their local tissue microenvironments, providing defense against pathogens and malignancies^6–10^. During primary infections, both antigen (Ag)-specific and bystander T cells are recruited to tissues, where some may differentiate into TRM, while secondary infections can induce *in situ* proliferation of resident Ag-specific TRM cells to rapidly contain local infection^11^. Notably, functional HIV-specific CD8^+^TRM have been detected in the rectal mucosa and lymphoid tissue of HIV controllers, even in the context of low levels of viral replication^12,13^.

Differentiation of memory precursors into CD8^+^TRM phenotypes has been linked to instructive signals within both inductive lymphoid tissues and peripheral tissue micro- environments, which underscores the presence of early TRM progenitors in circulation^7^. Among these signals, interleukin (IL)-15 and transforming growth factor-β (TGF-β) isoforms are critical factors regulating persistence of Ag-specific and bystander TRM in several tissues^7,8^. Indeed, sequential exposure of peripheral blood mononuclear cells (PBMC) to Ag or IL-15, followed by TGF-β, induces expression of CD69 and CD103 in CD8^+^T cells, two molecules associated to TRM phenotypes in some tissues^6–10^. These newly induced CD69^+^CD103^+^CD8^+^ T cells concomitantly express immune checkpoint (IC) molecules like PD-1 and CD39^14,15^, but can still rapidly produce IL-2 and IFNγ, and are indirectly linked to hepatitis B virus control^16^. Similarly, CD69^+^CD103^+^ TRM-like cells with expression of CD39 accumulate in various human solid cancers, as a consequence of hypoxia^17^ and/or recurrent TCR activation in a TGF-β-rich environment^18^, where they have been associated with improved disease outcome and patient survival, likely due to enrichment of Ag-specificity^19,20^. These phenotypes may comprise *bona fide* TRM and dysfunctional exhausted T cells that exhibit a variety of TRM- like features driven by continuous antigen exposure within the microenvironment^18,21,22^. In fact, chronic Ag stimulation may drive distinct or unconventional residency programs characterized by upregulation of IC, which may indeed restrain proliferation and activity ^18,23^. Still, phenotypically, *bona fide* CD8^+^TRM subsets, as well as exhausted TRM-like profiles will vary among tissues, adding complexity towards their identification^17,21–24^.

In this study, we hypothesized that stimulating cells from blood with cytokines inducing CD8^+^TRM signaling could remodel effector cells towards tissue-adapted subsets, more likely to control viral rebound in tissues upon treatment interruption. Sequential IL- 15/TGF-β1 treatment of PBMC recovered antiviral CD8^+^T cells not limited by expression of CD39 and PD-1. Effector CD8^+^T cell subsets induced by cytokine-stimulation displayed phenotypic and transcriptional hallmarks of residency, strongly associated with metabolic reprogramming modulating inhibitory checkpoint expression and mitochondrial oxidative stress. This treatment enhanced control of the reactivated HIV reservoir in virally-suppressed people and effector phenotypes characterized by CD103 or CD39 expression together with CD69 were the only CD8^+^T cells subsets capable of eliminating intact HIV-1 viruses in PWH. In a humanized murine ATI model, adoptive transfer of CD8⁺ T cells sequentially exposed to IL-15 and TGF-β1 conferred superior control of HIV rebound compared with IL-15 treatment alone, with responses observed in a subset of animals. These findings identify phenotypic features of effective CD8^+^T cells that can control persistently infected cells and define a novel therapeutic approach to improve CD8^+^T cell function for HIV cure strategies and beyond.

## Results

### Sequential IL-15/TGF-β1 stimulation induces proliferating CD103^+^ and CD39^+^ CD8^+^T cells and restores polyfunctional HIV-specific CD8^+^T cells

To induce a potentially tissue-adapted phenotype in circulating CD8⁺T cells, we stimulated PBMC from PWH with IL-15 for three days, followed by three days of TGF- β1, based on prior work from Pallet et al^16^. As shown for a representative sample (**Fig. 1a**), 2.94% of CD8⁺ T cells co-expressed CD69 and CD103 after treatment, a phenotype that was scarcely detectable at baseline or after six days of culture without cytokine stimulation. Among all tested PWH-samples, sequential treatment induced a significant increase of single CD69, CD103 and CD39 expression compared to baseline, without changes in PD-1 expression (**Fig. 1b**). The frequency of double CD69^+^CD103^+^ CD8^+^T cells was variable, with a median of 2.03% (IQR 1.21-5.19), which was significantly higher compared to baseline (**Fig. 1c**). Additional phenotypic combinations of interest in relation to TRM-like and Ag-enriched signatures during chronic antigenic stimulation^18,19,21,23,25^ were also significantly enhanced, namely CD69^+^CD39^+^, CD39^+^CD103^+^ and CD69^+^CD103^+^CD39^+^ CD8^+^T cells (**Fig. 1c**). Of note, compared to the CCR7^+^ fraction, effector CCR7^-^CD8^+^T cells contained most of these CD69^+^CD103^+^ and CD69^+^CD39^+^ subsets, as well as higher expression of Ki-67, Eomes and T-bet (**Extended Data Figs. 1a and b**). In fact, only CD69^+^ cells expressing CD39 or CD103 were enriched in proliferating Ki-67^+^ cells, since CD69^+^ only, or triple negative cells, were more frequent in the Ki-67^-^ fraction (**Extended Data Figs. 1a and c**). Among phenotypic combinations based on these markers, CD69^+^CD103^+^CD39^+^CD8^+^T cells showed the highest expression of Ki-67 and T-bet expression, while lower expression of Eomes was detected in this subset compared to CD69^+^CD103^-^CD39^+^ (**Extended Data Fig. 1d**).

**Fig. 1.**
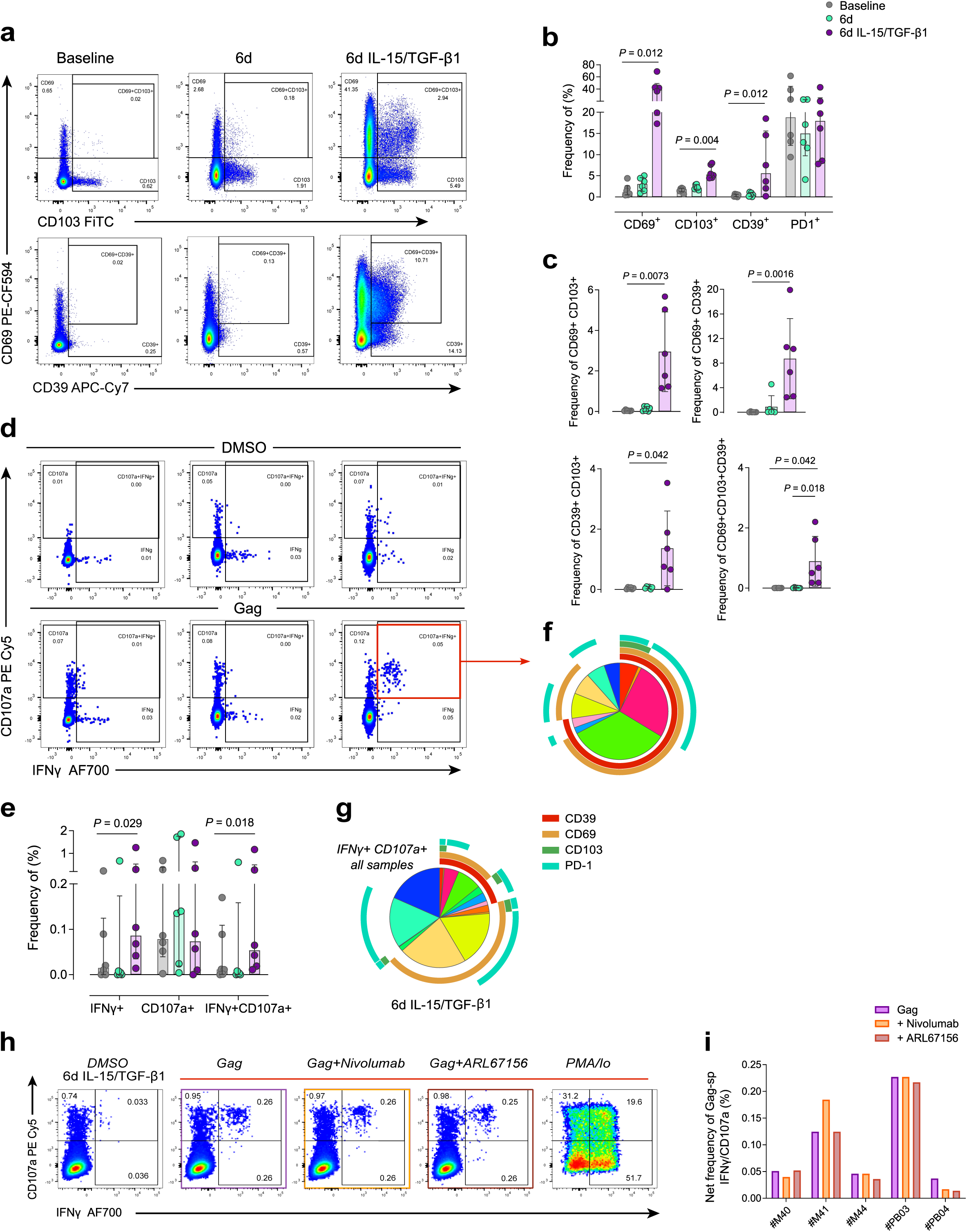
IL-15/TGF-β1-stimulation in PBMC from ART-suppressed patients enhances polyfunctional Gag-specific CD8^+^T cells not restricted by CD39 and PD-1 expression. a and d, Representative flow cytometry plots of baseline (left column), 6- days culture (center column) and 6-days culture with IL-15/TGF-β1 stimulation (right column) in a single patient showing the induction of CD8^+^T cells expressing CD69^+^CD103^+^/CD39^+^ (a) or CD107a/IFNγ in DMSO or Gag-stimulated conditions (d). b-c, Frequency of individual markers (b) or specified combinations (c) within total CD8^+^T cells at baseline (grey), after 6 days of culture (green) or after sequential cytokine treatment (purple) in ART-suppressed patients (n=6, from the Extended Data **Table 1**). e, Frequency of IFNγ and/or CD107a CD8+T cells in the same conditions and in the same ART-suppressed patients as in (b-c). Percentages represent net frequencies (DMSO subtracted) of IFNγ and/or CD107a Gag-specific blood CD8^+^T cells for each study conditions. For b, c and e, bars and error bars represent median and interquartile ranges. Statistical comparisons by one-way ANOVA with Dunn’s correction are indicated. f-g, Pie charts showing the proportions of expression of CD39, CD69, CD103 and PD-1 within the total double IFNγ^+^CD107a^+^ Gag-specific CD8^+^T cells determined by boolean gates after sequential treatment in (f) the example patient or (g) the average cells from all patients (n=6) after with cytokine-treatment. The colorful fractions inside each pie chart indicate the proportion of cells expressing a given phenotype indicated by the outside arches (see color legend for arches). h, Flow cytometry plots showing the frequency of IFNγ and CD107a Gag-specific CD8^+^T cells in PBMC from individual #PB03, stimulated with IL-15/TGF-β1 and further exposed to control (DMSO, left column) or stimulation with Gag (purple), Gag+anti-PD-1 (Nivolumab, orange), Gag+ATPase inhibitor (ARL67156, brown) or PMA/ionomycin (PMA/Io, right column). i, Net frequency (DMSO subtracted) of IFNγ^+^CD107a^+^ Gag-specific CD8^+^T cells in PBMC from five ART-suppressed patients stimulated with IL-15/TGF-β1 before further stimulation in the same condition as in (h). Statistical comparisons were performed using two-sided nonparametric Wilcoxon matched pairs signed rank test to compare the two groups.

Concurrently, a polyfunctional IFNγ/CD107a Gag-specific CD8^+^T cell response emerged only after treatment (**Fig. 1d**) and the net frequency (DMSO subtracted) of this response was statistically significant compared with baseline (**Fig. 1e**). These recovered anti-Gag IFNγ/CD107a CD8^+^T cells were composed of cells expressing CD69 (orange arch, **Figs. 1f and g**), with or without CD39 and PD-1, and low expression of CD103 (**Figs. 1f and g**). Noteworthy, this cytokine treatment did not induce Gag-specific responses in CD4^+^T cells or CD3^-^ lymphocytes with dim expression of CD8 (**Extended Data Fig. 2**). Since PD-1 and CD39, whose expression could indicate exhaustion^14,15,26,27^, were often represented among Gag-specific CD8^+^T cells after expansion, we tested the effect of blocking PD-1 or inhibiting CD39 activity during Gag stimulation with Nivolumab or with the ATPase inhibitor ARL67156, respectively ^28,29^. As shown (**Figs. 1h and i**), the net frequency of polyfunctional Gag-specific CD8^+^T cells was not modified for any of the five PWH-samples tested (**Fig. 1i**), except for one, which showed enhancement of IFNγ secretion when PD-1 was blocked. Of note, this participant had a long history of HIV infection and multiple co-infections, as well as repeated ART interruptions, collectively suggesting a more immunologically exhausted profile ^40, 42, 46^. These results demonstrate that treatment of PBMC with IL-15 followed by TGF-β1 induces proliferation of effector CD8^+^CD69^+^T cells expressing CD103 and/or CD39, while recovers a polyfunctional antiviral CD8^+^T cell response, a desired feature of effective HIV-specific CD8^+^T cells not fully restored by ART^30^. Considering that in most individuals this response is not limited by high expression of PD-1 or CD39, in the rest of the study we refer to CD39 expression after sequential treatment as part of the TRM- like phenotype (induced by TGF-β1 as reported^19^), rather than as an exhaustion marker.

### CD8^+^T cell induced by sequential IL-15/TGF-β1 expressing CD103 or CD39 display enhanced elimination of the viral reservoir

Considering the enhancement of HIV-specific CD8^+^T cells after sequential stimulation, we established a functional assay to compare the ability of treated CD8^+^T cells to eliminate the autologous HIV reservoir after viral reactivation.

CD4⁺T cell reactivation was induced pharmacologically with the latency-reversing agent (LRA) Ingenol^31^ and assessed by flow cytometry (**Fig. 2a**). Using samples from six PWH, we conducted two independent experimental series to compare the efficacy of IL- 15/TGF-β1-treated CD8⁺T cells with that of 1) untreated CD8^+^T cells (**Fig. 2b**-**d**), and 2) IL-15-treated CD8^+^T cells (**Fig. 2e**-**g**). We included IL-15 treatment as a control because it has been shown to reshape CD8^+^T cells and enhance cytotoxicity^32,33^ and clinical trials using IL-15 to promote post-ART control of HIV are ongoing (NCT04340596 and NCT07145164). Reactivation significantly induced p24 expression compared to baseline, which was significantly reduced only when CD4^+^T cells were co-cultured with 1:1 autologous IL-15/TGF-β1-treated CD8^+^T cells (**Fig. 2b**). Paired comparison of residual p24 frequencies revealed significantly lower levels in IL-15/TGF-β1-treated CD8^+^T cells compared with untreated controls (**Fig. 2c**). By normalizing to 100% the individual level of p24 per patient after reactivation, we observed that CD8^+^T cells cultured for six days without any treatment reduced p24 expression by a median of 33%, while IL-15/TGF-β1- treated CD8^+^T cells reduced p24 levels by a median of 75% (**Fig. 2d**). In the second set of samples from PWH, only sequential IL-15/TGF-β1 treatment, and not three days of IL-15 alone, significantly reduced the frequency of p24-expressing CD4^+^T cells after reactivation (**Fig. 2e**), resulting in significant differences between treatment strategies (**Fig. 2f**), although both conditions reduced the number of infected cells (**Fig. 2g**).

**Fig. 2.**
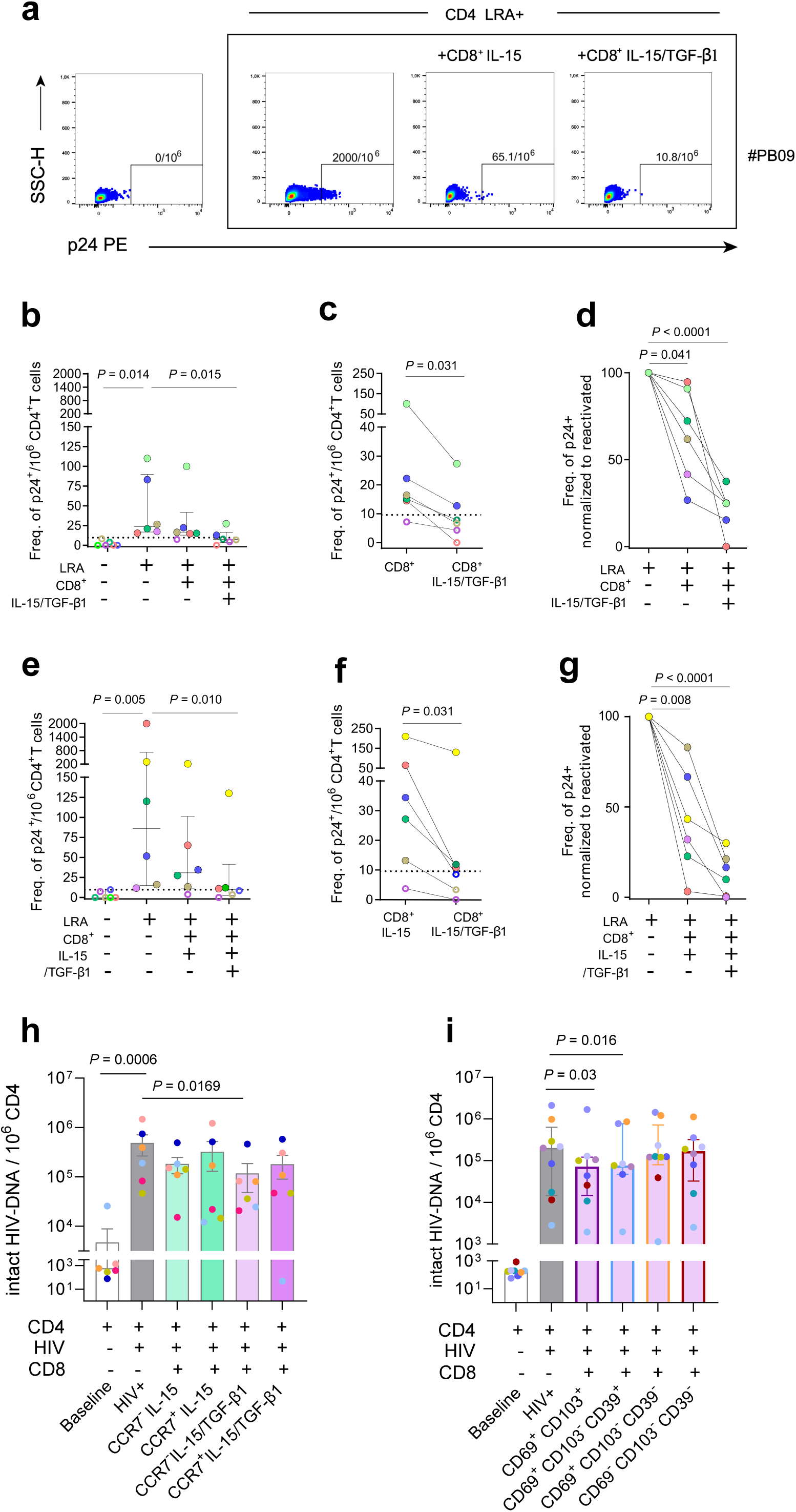
Enhanced killing of autologous reactivated CD4^+^T cells and intact proviruses from ART-suppressed people after sequential IL-15/TGF-β1 treatment. **a,** Flow cytometry plots of p24 detection in blood CD4^+^T cells from a PWH (#PB09) at baseline (not reactivated, left plot), after reactivation with Ingenol as a latency reversal agent (LRA) alone, or in the presence of CD8^+^T cells 1:1 that were stimulated with IL-15 only or with IL-15/TGF-β1 (right plot). The number of p24^+^ cells per million CD4^+^T cells is indicated for each plot. **b** and **e,** Number of p24^+^ cells per million CD4^+^T cells after co- culture of LRA-reactivated CD4^+^T cells with unstimulated (CD8, **b**) or with IL-15 stimulated (**e**) *versus* IL-15/TGF-β1 stimulated CD8^+^T cells. Open symbols are below the limit of detection (9.6 molecules/10^6^ CD4^+^T cells), which is indicated with a dotted line. Statistical comparisons were performed using two-sided non-parametric Friedman with Dunn’s multiple comparison test. **c-f**, Paired comparison of the detection of p24 expression per million CD4^+^T cells after the addition of untreated (**c**) or IL-15- treated (**f**) *versus* IL-15/TGF-β1-treated CD8^+^T cells, as in **b** and **e**. Statistical comparisons were performed using two-sided non-parametric Wilcoxon matched-pairs signed rank test. **d- g,** Killing capacity of untreated (**d**) or IL-15-treated (**g**) *versus* IL-15/TGF-β1-treated CD8^+^T cells normalized to the detection of p24 in reactivated-only CD4^+^T cells (as the 100%). Statistical significance was assessed using a two-sided non-parametric one-way ANOVA, followed by Dunnett’s multiple-comparisons test. **h-i**, Bar chart showing molecules of intact HIV-DNA per million CD4^+^T cells at baseline, or after *ex vivo* reinfection of CD4^+^T cells from PWH co-cultured, or not (in grey), in the presence of indicated CD8^+^T cell subsets obtained from: **(h)** stimulated IL-15 (in green) and IL- 15/TGF-β1-treated (in purple) PBMC and **(i)** from IL-15/TGF-β1-treated PBMC only, as shown in gating strategy (**Extended Data** Fig. 3c). Median lines and error bars represent median and interquartile ranges. Statistics were performed using two-sided nonparametric Wilcoxon matched-pairs signed rank test for group comparison. Each color represents an individual sample as indicated: **b** to **d** (n = 6, ), **e** to **g** (n = 6), **h** (n=6) and **I** (n=9; **Extended Data Table 1**).

To confirm the increased capacity to eliminate HIV-infected cells of IL-15/TGF- β1-treated CD8^+^T over IL-15 only treatment, and verify that the CCR7^-^ effector fraction was the main responsible for this effect, we superinfected isolated CD4^+^T cells, which were cocultured with multiple autologous CD8^+^T cell subsets to determine the capacity to eliminate cells containing intact HIV-DNA^31^. As shown, sequentially-treated CCR7^-^ CD8^+^T cells were the only fraction of cells capable of significantly reducing the levels of intact HIV-DNA compared to the superinfected condition (p=0.005 **Fig. 2h**). In relation to defective forms, no significant differences were observed (**Extended Data Fig. 3a**). Of note, while the effector CD8^+^T fraction from both stimulated conditions were enriched in PD-1 and costimulatory receptor 4-1BB expression compared to the CCR7^+^ fractions, suggesting enhanced responsiveness to Ag^32^, the IL-15-stimulated condition presented the highest expression levels, (**Extended Data Fig. 3b**).

We next sought to determine which specific subsets from the IL-15/TGF-β1- stimulated CCR7⁻CD8⁺ T cell population were capable of selectively eliminating HIV- infected cells harboring intact proviruses. Since these phenotypes were limited in frequency, and high number of cells was necessary to perform this killing assay, we could only assess a limited number of subsets. Thus, we prioritized the CD69^+^CD103^+^subset, as a more conventional TRM-like phenotype^7,8,23,24,34^, regardless of CD39 expression, followed by the CD69^+^CD103^-^CD39^+^ subset, as an Ag-enriched unconventional TRM- like/exhausted phenotype^15,18–20,23^. These subsets were compared to activated CD69^+^CD103^-^CD39^-^ and triple negative subsets obtained from the same PWH (**Extended Data Fig. 3c**). As shown, both fractions significantly reduced intact HIV-DNA, in contrast to CD69^+^CD103^-^CD39^-^ and triple negative phenotypes, which were not significantly different from the infected condition (**Fig. 2i**). Of note, no changes in defective forms were observed compared to the superinfected HIV^+^ condition (**Extended Data Fig. 3d**). Concordantly, and despite being less abundant in frequency, these two subsets were also significantly enriched for PD-1 and 4-1BB in comparison to CD103^-^/CD39^-^ phenotypes, again suggesting increased response to Ag^32^ (**Extended Data Fig. 3e**). Overall, these results support the concept that, beyond the expansion of Ag-specific T cells induced by sequential treatment, only induced CD69^+^ CD103^+^ and/or CD39^+^ effector phenotypes hold enhanced capacity to significantly eliminate intact HIV forms.

### Single cell RNA sequencing reveals increased clonotype diversity and expansion of cycling T_RM_-like with enhanced mitochondrial capacity and limited exhaustion

Compared with IL-15, sequential treatment increased effector CCR7⁻CD8⁺T cells in samples from PWH (**Extended Data Fig. 4a**), and the frequency of cells separately positive for, or co-expressing, CD49a and CXCR3 (**Extended Data Figs. 4b and c**). These changes are consistent with acquisition of a cytokine-driven TRM-like program, together with enhanced potential for homing to inflamed tissues and tissue persistence^6,24,34^. Within IL-15/TGF-β1-treated samples, CD69⁺CD103⁺ and CD69⁺CD39⁺ subsets showed the highest frequencies of CD49a^+^ and CD49a^+^ CXCR3^+^ cells compared with total CD8⁺T cells, while only CD69⁺CD103⁺CD8⁺T cells were enriched in CXCR3 expression compared to total CD8⁺T cells (**Extended Data Fig. 4d**).

To investigate the effects of sequential treatment on these expanded CCR7⁻CD8⁺T cells in greater detail, we performed single cell RNA sequencing (scRNAseq) by 10X Genomics 5′scRNA-seq, along with V region sequencing of TCRA and TCRB genes of T cells in multiple samples from a unique PWH, in which, in addition to blood, we had the opportunity of a large cervical tissue sample for comparison (#M47, **Extended Data Table 1**). To increase the frequency of Ag-responding cells, we included Gag during the three-day stimulation with IL-15, which further increased the proportion of double IFNγ^+^/CD107a^+^ Gag-specific CD8^+^T cells after TGF-β1 stimulation (**Extended Data Fig. 4e**). Thus, we analyzed data from CCR7^-^CD8^+^T cells (range 5,635-7,916 cells) derived from blood from this participant at baseline, after IL-15+Gag treatment and after sequential IL-15+Gag/TGF-β1 treatment; all of them after overnight Gag re-stimulation or DMSO, as the control (gating strategy in **Extended Data Fig. 5a**). Concurrently, to include transcriptional signatures of TRM from a tissue of HIV persistence^35^, we performed scRNA-seq of CD45^+^ lymphocytes isolated from the cervix of the same participant (5,500 cells; gating strategy in **Extended Data Fig. 5b**), which was also Gag- stimulated overnight; and from a cervix of an non-HIV woman, as the DMSO control (5,430 cells). Of note, as expected, CD8^+^T cells from the #M47-cervical sample and from sequentially-treated PBMC showed the highest expression of CD103 and CD39, as well as of PD-1 protein expression, compared to the other PBMC conditions (**Extended Data Fig. 5c**).

The comparison of the overall gene expression of selected groups of genes of interest showed remarkable differences among the eight samples studied (**Fig. 3a-d**). Genes related with self-renewal and stemness were enhanced in the IL-15 treated condition from blood (**Fig. 3a**), which also promoted mTORC1 pathway products (*HIF1A*)^33^, genes associated to cytotoxicity (**Fig. 3c**) and exhaustion (**Fig. 3d**). In contrast, sequential treatment induced expression of a variety of genes associated to tissue residency^7,10,12,36^, including genes associated to adhesion and cytoskeletal remodeling (*ITGA2, VIM*, *ANXA2, TUBB*), chemokine receptors (*CCR5, CXCR6*) and transcriptional factors (*ZNF683* -HOBIT-, *RBPJ*)^36–38^ (**Fig. 3b**). Consistent with protein expression patterns observed in previous samples, the genes encoding CD39 (*ENTPD1*), CD103 (*ITGAE*), CD49a (*ITGA1*) and CXCR3 were upregulated after sequential treatment, while *CD69* gene expression was downregulated compared to IL-15-only treatment (**Fig. 3b**). In addition, sequential treatment showed enhancement of transcriptional factors associated to memory (*EOMES*, *AHR*)^39^, reinvigoration of exhausted CD8^+^T cells (*KLF4, JUN*)^40,41^ and genes associated to mTORC2 pathway (*RICTOR*), a pathway linked to enhanced natural control of HIV-1^33^ (**Fig. 3a-b**). Of interest, the highest expression of some of these genes, including *KLF4, JUN, AHR* together with *ENTPD1,* was observed in the uninfected cervical control sample (**Fig. 3a-b**). In addition, following sequential stimulation, cells exhibited an overall less exhausted transcriptional profile, including reduced expression of inhibitory or terminal differentiation markers such as *KLRG1, LAG3, CTLA4*, *HAVCR2* (TIM-3), *PDCD1* (PD-1) or *CD160*^21,26,27^ (**Fig. 3d**). An exception was *CD101*, an inhibitory marker commonly associated with TRM-like signatures ^10,21,24^, which was upregulated in IL-15/TGF-β1-treated cells, with the highest expression observed in the cervical control condition. Last, sequential treatment increased proliferation-associated genes (*MKI67*, *PCNA*, *TOP2A*) and oxidative phosphorylation-related mitochondrial transcripts (*MT-CO1/3, MT-CYB*)^10^ in treated cells from blood (**Fig. 3d**).

**Fig. 3.**
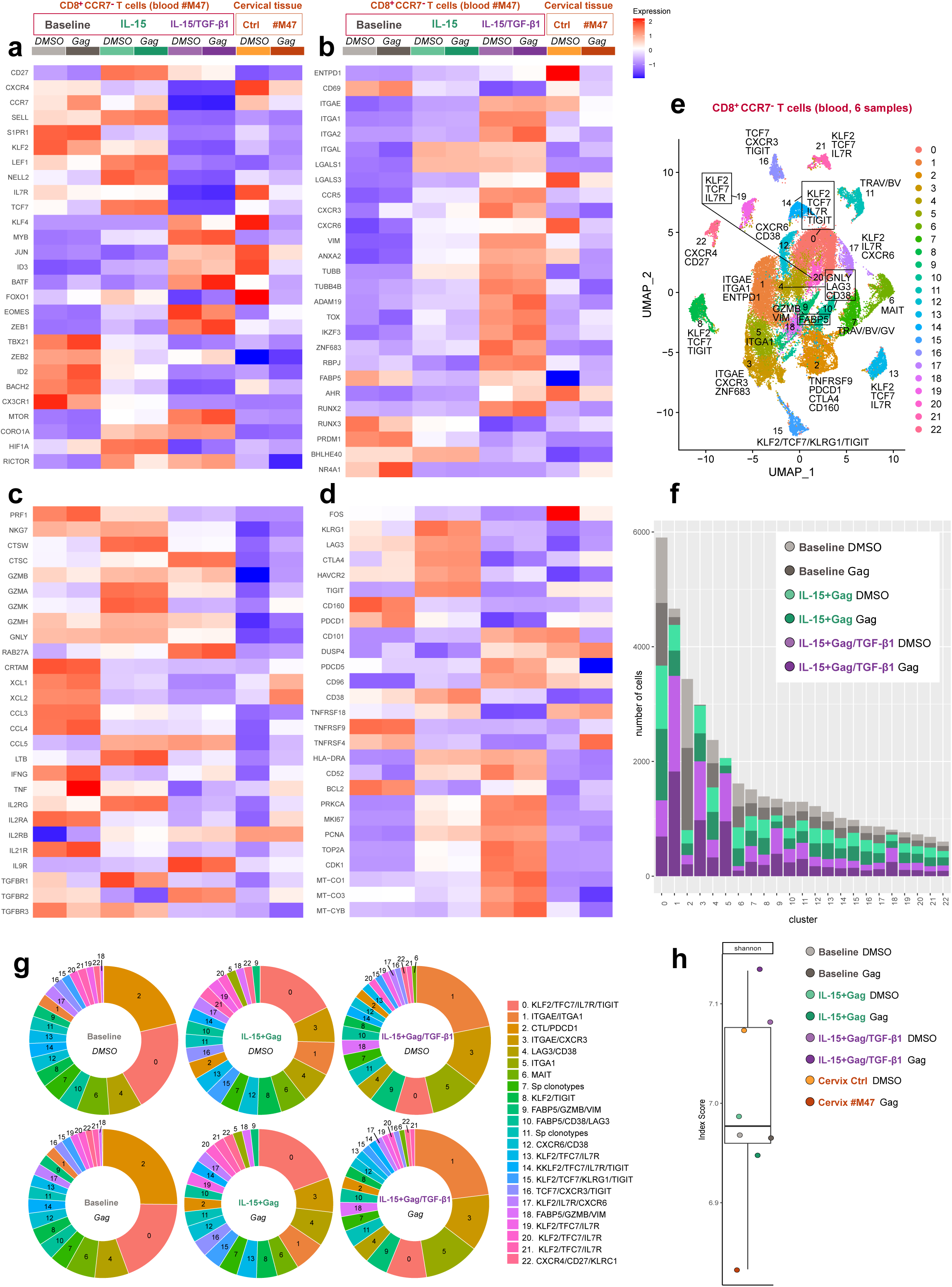
Single-cell transcriptome profiling of effector CD8^+^T cells and clonotype diversity after IL-15/TGF-β1-treatment. **a-d**, Heatmap displaying log-normalized average expression of selected markers annotated by functional groups for the eight conditions studied (as indicated in the top labeling). **e,** UMAP visualization of CCR7⁻CD8⁺ T-cell clusters identified at a resolution of 0.5 based on highly variable genes and integrated across PBMC samples from six conditions. **f,** Stacked bar chart showing cell number per cluster for each sample, based on the 22-clusters identified in PBMC as in (**e**). **g,** Pie charts showing relative cluster abundance for each blood sample after DMSO (top) or Gag (bottom) overnight stimulation. Legend indicates cluster number, highlighting 1 to 3 genes of interest upregulated (as indicated in the corresponding UMAP (**e**) for reference). “Sp clonotypes” denote clusters with few differentially expressed genes, distinguished primarily by the expression of specific TCR genes. **h**, Shannon index indicating repertoire diversity used to estimate the clonotypic richness of each sample, considering all eight samples as indicated in the color code legend.

We next focused on the analysis considering only the six PBMC conditions to identify clusters expressing *ITGAE* and *ENTPD1* genes after sequential treatment, as they correspond to CD103 and CD39 protein expression, phenotypes that we identified as the most effective at eliminating HIV. The clustering of highly variable genes across PBMC samples identified 22 distinct clusters (**Fig. 3e**). Since cell sorting was based on the lack of CCR7, these clusters were mostly composed of effector CD8^+^T cell populations, including MAIT cells, identified based on genes like *TRAV1-2, SLC4A10 and KLRB1*^10^, as cluster 6 (**Fig. 3e**). At baseline conditions, clusters 2 and 0 were the most abundant (**Fig. 3f-g**). Cluster 0 exhibited a transcriptional program associated with self-renewal and quiescence, consistent with memory precursor cells^7^ (*KLF2, TCF7*, *IL7R, SELL*), which co-expressed *TIGIT*, a profile shared with several minor clusters (8, 13-16, 19 and 21) (**Fig. 3e**). In contrast, cluster 2 corresponded to terminally differentiated or short-lived effector cytotoxic T lymphocytes (SLECs), characterized by the expression of *TBX21, PRDM1, ID2* and *ZEB2*^7,36,37^ along with genes associated with activation, cytotoxicity and exhaustion (*CD69, TNFRSF9, IFNG, PRF1, GZMB, PDCD1, CD160, LAG3, CTLA4, HAVCR2*). Besides cluster 2, additional clusters (4, 6, 9, 10, 12, and 18) also showed elevated expression of cytotoxic genes.

Sequential IL-15/TGF-β1 treatment induced expansion of clusters 1, 3, and 5 (**Fig. 3f and g**), which were related by trajectory (**Extended Data Fig. 6a**), and to a lesser extent of clusters 9 and 18, while reducing the abundance of others, including clusters 0, 2 and 6 (MAIT) (**Fig. 3f and g**). Cluster 1, the dominant population, was uniquely characterized by the co-expression of *ITGAE* (CD103, also expressed in cluster 3) and *ITGA1* (also expressed in cluster 5), together with exclusive expression of *ENTPD1* (CD39). This cluster was strongly associated with an increase in oxidative phosphorylation-associated genes (*MT-CO1/3, MT-CYB*)^10^, survival (*BCL2* and *PRKCA*)^33^, TRM signatures (*RUNX3, RUNX2, IKZ3, RBPJ*)^7,37,42^ and exclusive expression of *TGFBR2*, associated to TRM generation^43^. Of note, Gene Ontology analysis revealed a significant enrichment of terms associated with epigenetic modifications in this cluster (**Extended Data Fig. 6b**). Cluster 3, expressing *ITGAE* together with genes such as *CXCR3, ZNF683* and *FABP5*, cluster 5 (*ITGA1* and *RBPJ*) and cluster 9 shared expression of proliferation-associated genes (*MKI67*, *TOP2A*, *CDK1*). These clusters may align with ‘cycling TRM-like’^10^ and were more abundant on g2M and S phases (**Extended Data Fig. 6c**). In addition, these expanded clusters showed enrichment of the mTORC2 pathway^33^, since clusters 1 and 5 expressed *RICTOR* and clusters 3 and 18 expressed *RHOA*. Moreover, clusters 5 and 18 expressed *MYB*, which induces long-term preservation of self**-**renewal capacity in exhausted T cells^44^; while clusters 3 and 5 expressed *JUN*, shown to promote exhaustion resistance over the expression of *FOS* (the latter expressed in clusters 2 and 6)^40^. As expected, IL-15-only treatment generated and in between situation, and most abundant clusters were 0, 3, 1 and 4, this last one related to an IFN-stimulated gene signature (*STAT1, IFI6, ISG15*) (**Fig. 3g**).

To address if clusters expanded after sequential IL-15/TGF-β1 treatment originated from few hyper expanded clones or not, we assessed their clonality, which indicated that they were mostly composed of individual or small clones (1 to 5 clones; **Extended Data Fig. 5c**). Actually, except for clusters 2 (putative SLECs), 7, 15 and 16, which showed greater proportion of large or medium clones, all other clusters showed predominance of single or small clones (≥ 65% of the cells; **Extended Data Fig. 6d**). Further, an overall estimation of the richness of the samples, measured by the Shannon Index, exposed higher diversity in sequentially treated samples compared to the other conditions, including the corresponding #M47 cervical sample (Gag), which showed the most restricted clonotypic repertoire (**Fig. 3h**). Of note, baseline and IL-15-only treatment showed a decrease in diversity index after Gag stimulation compared to the DMSO control, while sequential treatment displayed enriched diversity not only in the DMSO sample, but even higher after Gag stimulation (**Fig. 3h**).

Last, we considered all samples (6 from PBMC and 2 from cervix), which also identified 22 clusters (**Extended Data Fig. 7a**), to determine specific features of putative Ag-specific clusters. Based on the expression of genes like *TNFRSF9, IFNG* and *GZMB*, clusters 5, 7, 8 and 13 were identified as potentially Gag-responding CD8^+^T cells (**Extended Data Fig. 7b**). Of note, these clusters were related by trajectory (**Extended Data Fig. 8a**) and clusters 5 and 7 showed the highest clonal overlap (**Extended Data Fig. 8b)**. In samples from sequential treatment, these clusters exhibited enhanced expression of tissue-residency features, including the absence of *KLF2* and the expression of *ITGAE* and *ITGA1* (**Extended Data Fig. 7b**), while showed significant clonal overlap with clusters 2 and 3 (**Extended Data Fig. 8b**), corresponding to the expanded clusters after sequential treatment (**Extended Data Fig. 8a**). Since cluster 7 was the only cluster with upregulation of genes such as *TNFRSF9, IFNG* and *GZMB* in all Gag conditions, including cervix (**Extended Data Fig. 7b**), we performed differential expression analysis (DEA) of cells from this cluster among conditions (**Fig. 4a-d**). Compared to Gag- stimulated cervical CCR7^-^CD8^+^T cells, Gag-stimulated cells from blood within this cluster showed upregulation of *LAG3, HAVCR2, CD160* and *CTLA4* at baseline (**Fig. 4a**); of *LAG3, HAVCR2, KLRG1* and *TIGIT* after IL-15 treatment (**Fig. 4b**), and only of *LAG3* after sequential treatment (**Fig. 4c**). Other genes upregulated in IL-15/TGF-β1 treated cells relative to cells from the cervix included oxidative phosphorylation (*MT-C01, MT- CO3*) and proliferation-associated genes (*MKI67*, *TOP2A*), among others (**Fig. 4c**). Further, the comparison between IL-15 only and sequentially-treated cells indicated that, subsequent exposition to TGF-β1 in this sample induced upregulation of genes related to residency (*ITGA1, ITGAE, RUNX2, RBPJ*), oxidative phosphorylation (*MT-CO3*), proliferation (*MKI67*), function and survival (*IFNG* and *BCL2*) and downregulation of the inhibitory molecules (*LAG3, KLRG1, CTLA4, HAVCR2*), which were overexpressed within IL-15-only-treated cells from this cluster together with molecules associated to exhausted trajectories in tissue microenvironments such as *TIGIT, GZMA, GZMK, CD38, BHLHE40* or *FOS,* ^21,22,40^ (**Fig. 4d**). These differential patterns were partially retained in the DEA performed considering all clusters (**Fig. 4e-h**), Overall, these results indicate that, in this individual, sequential treatment increased clonotypic diversity in effector CD8^+^T cells, expanding transcriptional phenotypes associated to tissue residency with enhanced proliferation, mitochondrial function and downregulation of exhaustion hallmarks.

**Fig. 4.**
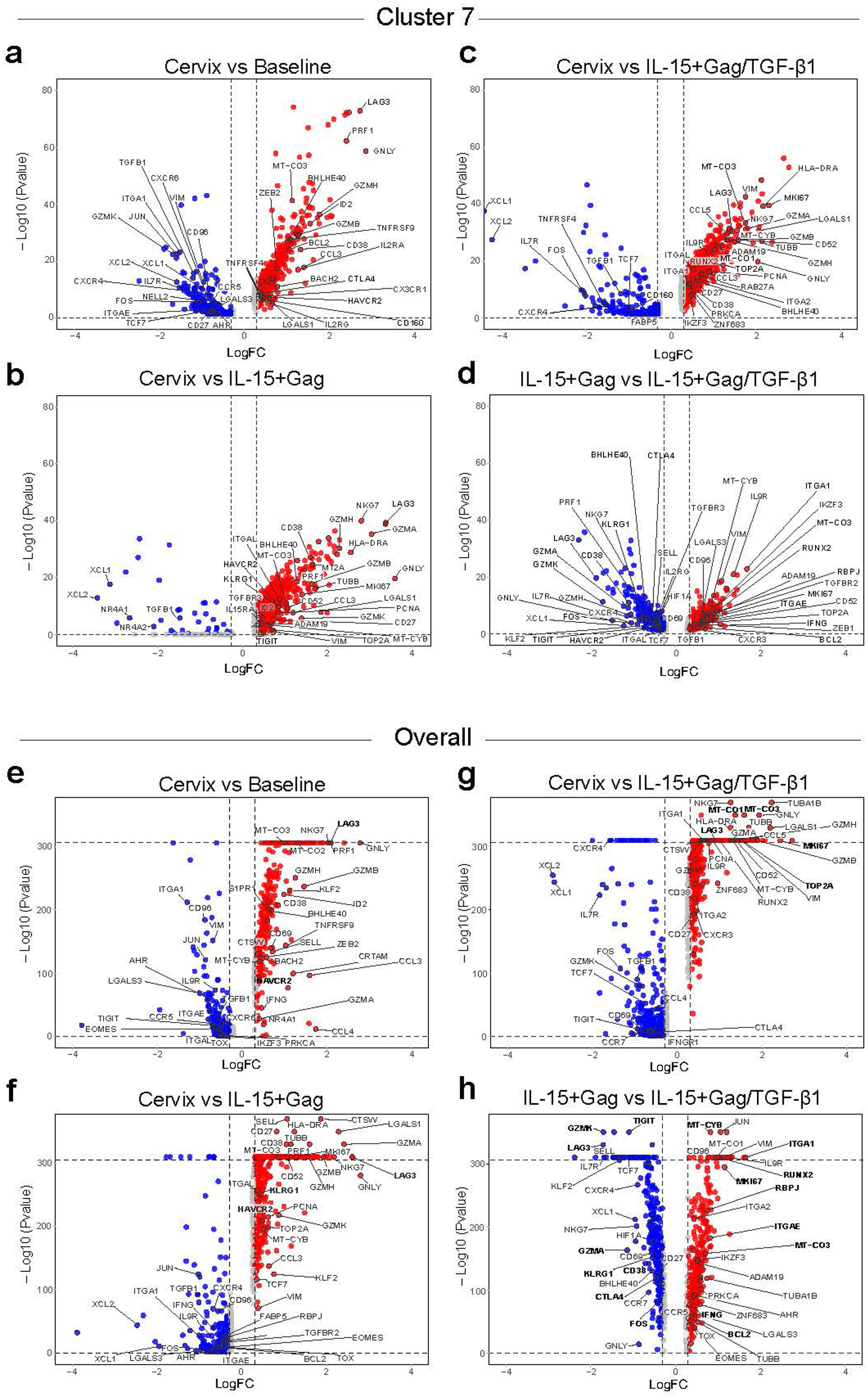
Differential expression analysis in a cluster of Ag-responding or in total CCR7^-^CD8^+^T cells among PBMC and tissue samples. **a**-**h**, Volcano plots showing differential expression analyses (DEA) considering (**a-d**) cluster 7 or (**e-h**) all clusters (overall) between (**a-c** and **e-f**) the cervical sample (#M47 Gag) and the matching Gag- stimulated PBMC condition or (**d** and **h**) between IL-15-only and IL-15/TGF-β1 treated PBMC (after Gag stimulation). Significantly upregulated genes (*P* < 0.05, log2-fold change (FC) > 0.3) are depicted in red, and significantly downregulated genes (*P* < 0.05, log2-FC < – 0.3) are depicted in blue. Top dotted line in the Y axes indicate a -Log10 (p value) of above 310 (limit of detection). Selected genes of interest are in bold.

### Sequential IL-15/TGF-β1 stimulation diversifies effector CD8^+^T cells subsets increasing intermediate phenotypes identified in tissue

To validate findings from transcriptional data, based on a single individual, we used spectral flow cytometry to assess the simultaneous expression of molecules associated to hits related to cell differentiation/homeostasis (CD27, IL-7R, CD52), residency/Ag-TRM- like (CD69, CD103, CD49a, CD39), exhaustion/terminal differentiation (CTLA4, LAG- 3, TIGIT, PD-1, KLRG1) and function (GzmB and IFNγ) in a new set of eight PBMC samples from PWH, including four with concurrent cervical biopsies (**Extended Data Table 1**). The expression profile of these molecules in effector CCR7^-^CD8^+^T cell subsets was analyzed under the same conditions as for the scRNA-seq. Despite substantial inter-sample variability, these analyses confirmed that residency markers were elevated after sequential treatment as well as in cervical samples (**Fig. 5a**), while KLRG1 was significantly higher in blood at baseline compared to the rest of the samples (**Fig. 5a**). Additional differences, when comparing all samples, were detected for IL-7R, downregulated after IL-15 compared to baseline in blood; and for CD52, upregulated in cervical samples compared to baseline (**Fig. 5a**). Notably, paired comparison between IL- 15 and sequential treatment indicated lower expression of TIGIT (*P* = 0.0078) and increased expression of IL-7R and CD27 together with PD-1 after IL-15+Gag/TGF-β1 stimulation (*P* = 0.0391; **Fig. 5b**). We also determined IFNγ responses and assessed their phenotype, as shown for an example patient (the same in which scRNAseq was performed, #M47, **Fig. 5c** and **5d**). Overall, higher IFNγ^+^ secretion was confirmed after sequential treatment and was significantly increased compared with both baseline and IL- 15-only treatment in paired analyses (*P* = 0.0078 and *P* = 0.0156 respectively, **Fig. 5e**). Moreover, compared to baseline, or after IL-15 treatment, the phenotype of IFNγ- secreting cells from blood following IL-15+Gag/TGF-β1 stimulation more closely resembled the one observed in cervix (**Extended data Fig. 9**).

**Fig. 5.**
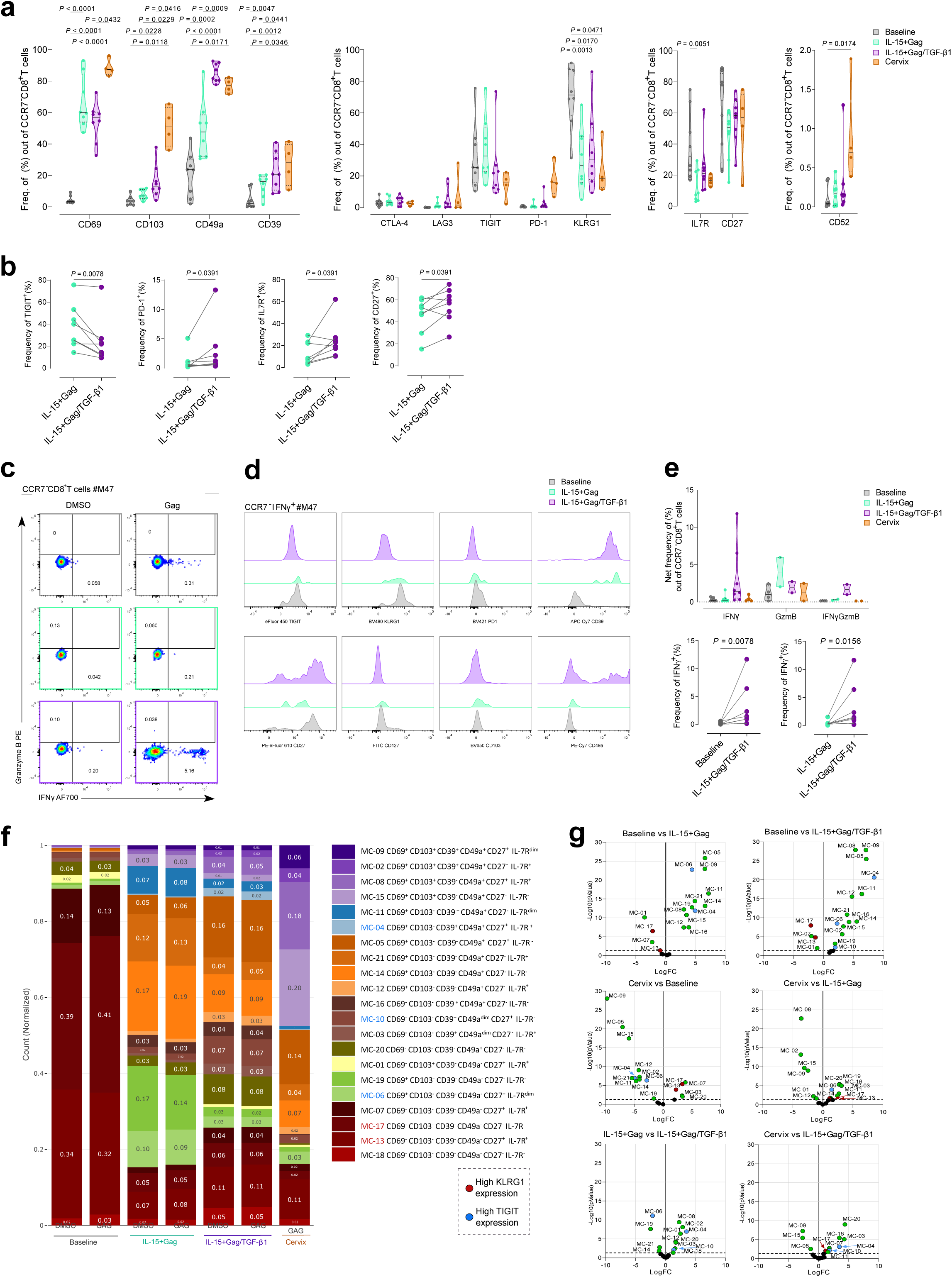
Phenotype of effector CD8^+^T subpopulations induced by sequential cytokine- treatment compared to blood and cervical samples. **a**, Protein expression of selected molecules in CCR7^-^CD8^+^T cells from PBMC-conditions from PWH (n=8; **Extended Data Table 1**) at the three study conditions (Baseline in grey, IL-15+Gag in green and IL-15+Gag/TGF-β1 in purple) and in four matched cervical samples (in orange). **b,** Paired comparison of the expression of different molecules within CCR7^-^CD8^+^T cells between indicated conditions. **c,** Flow-cytometry plots showing the expression of IFNγ and Granzyme B^+^ (GzmB) in Gag stimulated condition and corresponding DMSO control at baseline Baseline (top), IL-15+Gag (middle) or after IL-15+Gag/TGF-β1 treatment (bottom) for an example participant (#M47; **Extended Data Table 1**). **d,** Stacked overlay histograms showing the expression of TIGIT, KLRG1, PD-1, CD39, CD27, IL-7R, CD103 and CD49a in study PBMC-conditions within CCR7^-^IFNγ^+^ CD8^+^T cells of the same participant (#M47). **e**, Net frequency (top, DMSO subtracted) of Gag-specific IFNγ^+^ and GzmB ^+^ within CCR7^-^CD8^+^T cells from blood and cervical samples. Paired comparison (bottom) of the expression of IFNγ within CCR7^-^CD8^+^T cells between IL- 15+Gag/TGF-β1-expanded cells and the other two PBMC conditions. Statistical comparisons (**a**) and (**e**, top) were performed using a mixed-effect model and Tukey’s multiple comparisons test for multiple comparisons while for (**b**) and (**e**, bottom), Wilcoxon matched-pairs signed rank test were performed. Data is shown as individual samples, while lines represent median values and dotted lines interquartile ranges for (**a**) and (**e**, top). **f,** Stacked bar chart showing the normalized abundance (counts) of each identified cluster along the three PBMC-conditions and cervix within CCR7^-^CD8^+^ T cells, after DMSO or Gag-stimulation condition. Clusters were color-scale grouped by the expression of CD69, CD103, CD49a, CD39, IL-7R and CD27. **g**, Volcano plots showing differential cluster abundance between the indicated sample groups, as determined by two-sided edgeR analysis. Green dots indicate statistically significant differences (P < 0.05). Statistically significant clusters with high KLRG1 expression are shown in red, whereas those with high TIGIT expression are shown in blue in **f** and **g**.

Additional unsupervised clustering analysis of these samples, based on the mean fluorescence intensity of CD69, CD103, CD39, CD49a, CD27 and IL-7R, identified 21 clusters (**Fig. 5f and Extended data Figs. 10a and b**). Baseline conditions were dominated by clusters C17 and C13, both expressing the highest KLRG1 levels of all clusters (**Fig. 5f**), significantly enriched at baseline compared to all other samples (**Fig. 5g**). IL-15-treated samples expanded several clusters compared to baseline, including CD27^-^ IL-7R^-^ CD69^+^ expressing C19 and C14 clusters, but also phenotypes characterized by TIGIT expression such as C06 (**Fig. 5f**), which was enriched in this condition compared to all other samples (**Fig. 5g**). Sequential treatment further increased the number of clusters detected, overall increasing clusters that together with CD103 and/or CD39 expressed CD27 and/or IL-7R, such as C02, C04 and C08 (**Fig. 5f)**. Cluster C02 expressed the highest levels of CD103, together with CD39, IL-7R and IFNγ, and this cluster was significantly upregulated after sequential treatment compared to the other blood samples, but not to cervix (**Fig. 5g**). Indeed, compared to the cervical sample, sequential treatment resulted in fewer and weaker differences than the other blood conditions (**Fig. 5g**). Altogether, these analyses suggest that sequential treatment expands less-differentiated intermediate effector phenotypes characterized by CD27 and IL-7R expression in combination with CD103, CD39 and/or CD49a expression, suggesting development of memory cells among effector pehnotypes^45^. Overall, these phenotypes are more competent at secreting IFNγ in response to Ag and are associated to decreased KLRG1 and TIGIT expression compared to baseline and IL-15-only treatment, validating scRNAseq data.

### Sequential IL-15/TGF-β1 stimulation enhances mitochondrial function of CD39 expressing cells

To further validate transcriptional data, we next evaluated mitochondrial function after overnight Gag re-stimulation or DMSO in three additional PWH. We determined the content of mitochondrial reactive oxygen species (ROS) by evaluating the proportions of cells with high levels of MitoSox-probe staining, as a readout of mitochondrial dysfunction and/or hyperactivation. As a control of mitochondrial content and size, we also used MitoTracker staining in parallel. The proportion of high MitoSox, out of the CCR7^-^CD8^+^T cell fraction (**Fig.6a**), which tended to increase in both DMSO and Gag- stimulated conditions after IL-15+Gag treatment compared to baseline, significantly decreased after sequential stimulation with TGF-β1 (**Figs. 6a and b**). Then, we compared the expression of CD39, CD69, CD103 and PD-1 in total and in MitoSox^high^ effector CD8^+^T cells (**Fig. 6c**). While CD39 and PD-1 were enriched in MitoSox^high^ cells from baseline and IL-15 conditions, sequential treatment only maintained significantly higher PD-1 expression in this fraction (**Fig. 6d**). Thus, although overall CD39 expression was induced by IL-15+Gag/TGF-β1 treatment, CD39-expressing cells better preserved mitochondrial function after sequential treatment, as they were significantly less frequent in the MitoSox^high^ fraction compared with baseline (P=0.0429; **Fig. 6d**). No differences were observed for CD103 or CD69 (**Fig. 6e**). Last, although out of the sequential treatment condition, no significant differences were detected among studied subsets (**Figs. 6 f and g**), CD69^+^CD103^+^ cells displayed in general the lowest proportion of MitoSox^high^ cells, while CD69^+^CD103^-^CD39^+^ cells showed the highest proportion which, as mentioned, gradually decreased based on cytokine treatment (**Figs. 6g and h**). Overall, the improved mitochondrial fitness of effector CD8⁺T cells after sequential treatment underscores the benefits of this treatment across multiple effector subsets, particularly reprogramming CD39^+^ exhausted phenotypes.

**Fig. 6.**
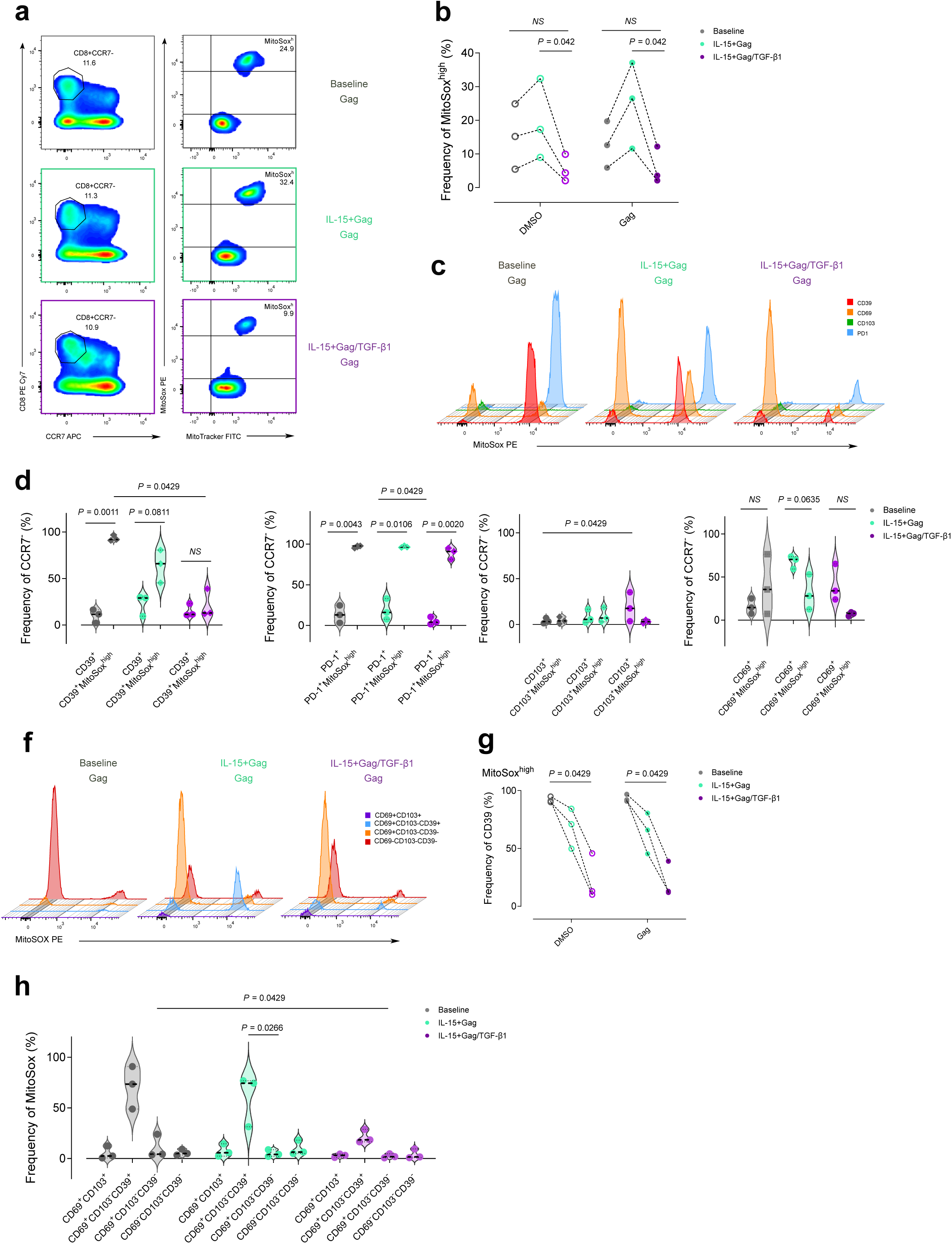
IL-15 /TGF-β1-treatment decreases mitochondrial reactive oxygen species in effector CD8^+^T cells. **a**, Flow cytometry plots showing CD8^+^CCR7^-^T cells (left panels) and of MitoTracker^+^ MitoSox^high^ (right panels) from a single individual (#PB07) after overnight Gag stimulation at: baseline (top, grey), IL-15+Gag (center, green) and sequential IL-15+Gag/TGF-β1 stimulation (bottom, purple). **b**, Frequency of MitoSox^high^ in CCR7^-^CD8^+^T cells after overnight DMSO or Gag stimulation in the three study conditions in PWH samples (n=3; **Extended Data Table 1**). **c**, Stacked overlaid histograms from the same individual sample (#PB07) showing the expression of MitoSox in CD39, CD69, CD103 and PD-1 expressing CCR7^-^CD8^+^T cells after overnight Gag stimulation in the three study conditions. **d-e**, Violin plots showing the percentage of expression of CD39 and PD-1 (**d**) or CD103 and CD69 (**e**) in total or in MitoSox^high^ CCR7^-^CD8^+^T cells after overnight Gag stimulation in the three study conditions from the same three patients as in (**b**). **f**, Stacked overlaid histograms in PBMC from the same individual (#PB07) showing the expression of MitoSox in CD69^+^CD103^+^ (purple), CD69^+^CD103^-^CD39^+^ (blue), CD69^+^CD103^-^CD39^-^ (orange), CD69^-^CD103^-^CD39^-^ (red) CCR7^-^CD8^+^T cells after overnight Gag stimulation in the three study conditions. **g**, Frequency of CD39 in MitoSox^high^ CCR7^-^ CD8^+^T cells after overnight DMSO or Gag stimulation in the three study conditions from the same patients. **h**, Violin plot showing the percentage of expression of MitoSox^high^ in the same subsets as in (**f**) after overnight Gag stimulation in the three study conditions from the same three patients. Dots represent individual samples and dotted lines medians. Open dots represent DMSO control samples while standard dots Gag-stimulated samples. Statistical analysis was done using non- parametric Friedman test for repeated measures with post hoc Dunn’s multiple comparisons test.

### Enhanced elimination of reactivated HIV^+^-CD4^+^T cells by sequentially- treated CD8^+^T cells in a humanized mouse model

Last, as a proof-of-concept, we evaluated sequentially-treated CD8^+^T cells as an immunotherapy to eliminate HIV reactivated cells from ART-suppressed PWH, by adapting humanized NOD-SCID 2Rγ–/– (NSG) mouse models^46,47^ to simulate ATI (**Fig. 7a**). For each PWH donor, sequentially treated CD8^+^T cells were compared to IL-15- treated CD8^+^T cells alongside a control group without CD8^+^T cells. This way, a total of 44 animals were allocated to three groups (controls n=19, IL-15-treated CD8⁺ T-cell recipients n=11, and IL-15/TGF-β1-treated CD8⁺ T-cell recipients n=14; **Extended data Table 2**). PBMC were obtained from eight PWH (**Extended data Table 1**), with one to four mice per condition per donor, based on cell recovery (**Extended data Table 2**). One day after irradiation, mice were intravenous (IV) injected with the same number of purified reactivated CD4^+^T cells, based on the PWH-donor, together with the remaining PBMC fraction (depleted from NK and CD8^+^T cells, **Extended data Table 2**). 5h after, animals received autologous CD8^+^T cells treated with IL-15 or IL-15/TGF-β1 (**Extended data Table 2**), except the control group, which remained without CD8^+^T cell adoptive transfer (**Fig. 7a**).

**Fig. 7.**
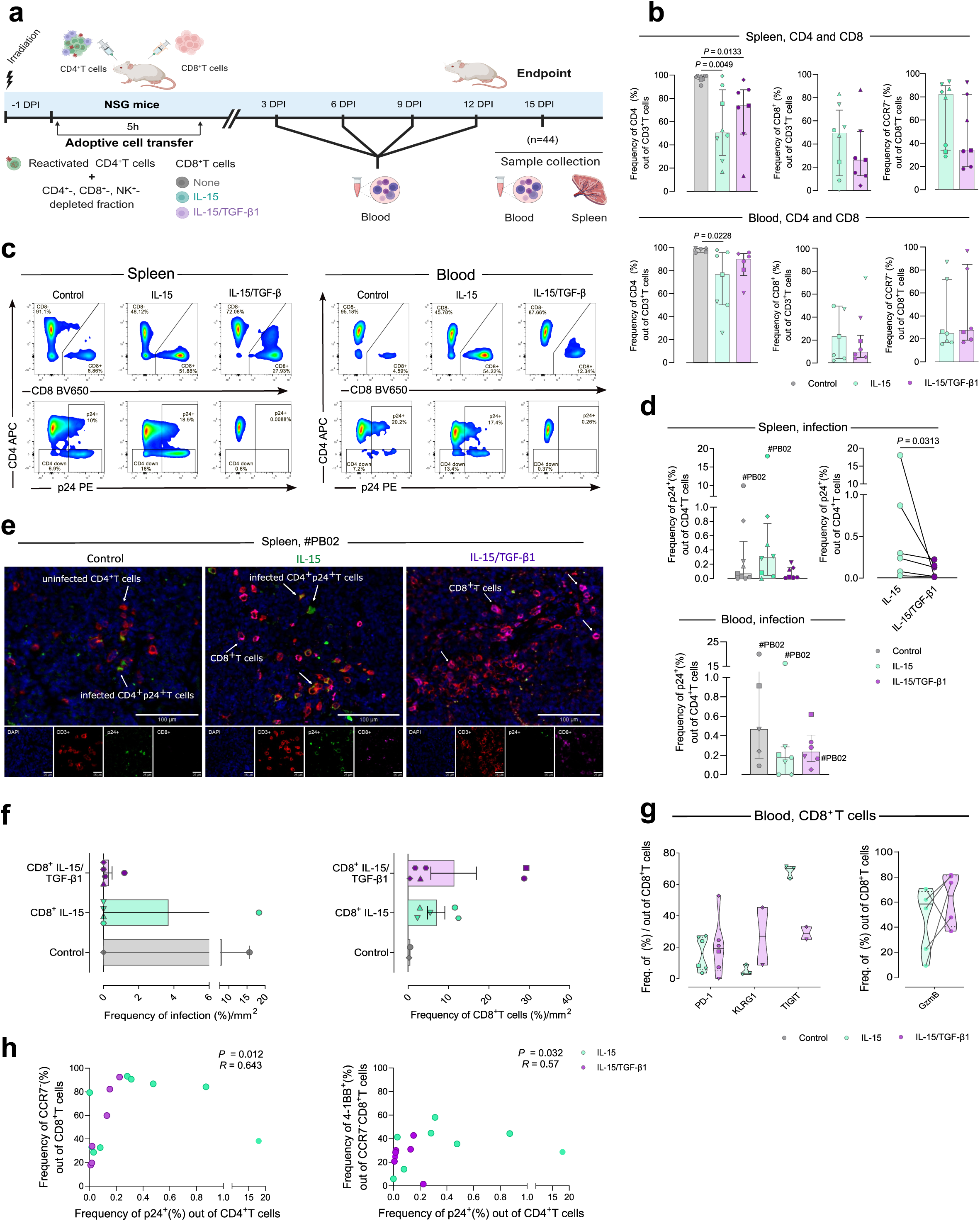
Evaluation of sequentially-treated CD8^+^T cells adoptive transfer to control viral reactivation in a humanized mouse model. **a,** Schematic representation of the experimental design. **b**, Bar charts showing the frequencies of CD4^+^T cells and CD8^+^T cells out of CD3^+^T cells or of CCR7^-^ cells in spleen (top) and blood (bottom) samples from the control (in grey), IL-15 (in green) and IL-15/TGF-β1 (in purple) mice groups at study endpoint. **c**, Flow cytometry plots from three independent mice belonging to each study group and derived from the same patient (#PB02, **Extended Data Table 1**) displaying the frequency of CD4 and CD8 in total CD3^+^T cells (top) or of p24 in total CD3^+^CD8^-^T cells (bottom) in samples from spleen (left) and blood (right). **d**, Bar charts showing the frequency of p24^+^T cells out of CD4^+^T cells from spleen (top) and blood (bottom) samples in the three study groups. In the spleen panel (top), the paired comparison between CD8-treataed conditions, performed by calculating the mean of the p24^+^ frequencies of mice belonging to the same group and patient, compared to the mean of the other group from the same patient, is shown. **e**, Microphotograph of immunofluorescence (IF) analysis showing anti-CD3 (red), anti-p24 (green) and anti- CD8 (pink) in splenic tissue of a control (left), IL-15 (middle) and sequential treatment mouse (right), corresponding to the #PB02 individual. White arrows indicate infected CD4^+^T cells (green and red), uninfected ones (red only) and CD8^+^T cells (pink). Bottom panels show magnified views of the indicated samples acquired at 20× magnification in multiple fluorescence channels. **f**, Horizontal bar charts from IF analysis of splenic frequencies of infection (p24^+^ cells) out of total CD3^+^CD8^-^T cells (left) and of CD8^+^T cells out of total CD3^+^ T cells (right) normalized by surface (mm^2^ of tissue) in detectable samples control (n=2), IL-15 (n=5) and sequential treatment mice (n=6) from six patients (**Extended Data Table 1 and 2**). **g**, Violin plots showing the expression of different molecules out of total CD8^+^T cells in blood of mice from the IL-15 (in green) or the sequential treatment group (in purple) and paired comparison of GzmB expression performed by calculating the mean of the frequencies of the mice belonging to the IL-15- group and patient, compared to the mean of the IL-15/TGF-β1-group and patient in samples with enough cell counts. **h**, Spearman correlation illustrating the association between splenic p24 expression in CD4⁺ T cells and CCR7⁻CD8⁺T cells (left) or with 4- 1BB⁺CCR7⁻CD8⁺T cells (left). For all flow cytometry spleen determinations, a total of 9 mice samples from the control group, 8 from IL-15 group and 7 from sequential treatment group were used from 6 patients (**Extended Data Table 1**). For flow cytometry blood determinations, a total of 5 mice samples from control, 7 from IL-15 and 6 from the sequential treatment group were used from 5 patients (**Extended Data Table 1**), except for KLRG1 and TIGIT expression with 2 samples from each mice group derived from 2 patients (#PB18 and PB20). All mice samples corresponding to each patient’s mouse group are indicated. Statistical comparisons in bar charts were performed using non- parametric Kruskal-Wallis test with post hoc Dunn’s correction for multiple comparisons while paired comparisons were analysed by Wilcoxon matched-pairs signed rank test. Data are shown as individual samples while bars represent median values and error bars interquartile range.

After T cell adoptive transfer, we were unable to detect HIV-RNA in plasma, probably due to low numbers of CD4^+^T cells in the low volumes obtained. We thus focused on the study endpoint, where only animals with enough cells were considered for the analyses (**Extended data Table 2**). As expected, the control group showed significantly higher frequencies of CD4^+^T cells out of total CD3^+^T cells in spleen compared to groups with CD8 adoptive transfer or, in blood, only compared to IL-15 treated mice (**Fig. 7b, left**). Also, no significant differences were observed in relation to the frequencies of CD8^+^T cells or the percentage of CCR7^-^ within this fraction between CD8-treated groups (**Fig. 7b**), however we noticed that spleens from the IL-15 group more frequently displayed high proportion of CCR7^-^CD8^+^T cells compared to the spleens from sequentially treated mice (**Fig. 7b, top right**).

The frequency of the viral protein p24 in CD4^+^T cells at endpoint in the three conditions from the donor #PB02, which showed the highest number of cells engrafted, indicated a strong reduction of reactivated p24^+^ CD4^+^T cells in the IL-15/TGF-β1-treated recipient mouse in spleen (from 10% to 0.009%; **Fig. 7c**, **top**) and blood (from 20.2% to 0.26%; **Fig. 7c**, **bottom**) compared to both controls. In these samples, high infection in control and IL-15 sample were accompanied by the downregulation of CD4 expression, a common feature of productively HIV infected CD4^+^T cells^35^ (**Fig. 7c**, **bottom row**). In samples from this donor we were able to perform a longitudinal analysis of the transferred T cell subsets, which in most time points showed > 100 CD45^+^ live cell counts, along with the frequency of p24^+^CD4^+^T cells (**Extended Data Fig. 11a**). This analysis confirmed a reduction in p24⁺ cells in the IL-15/TGF-β1-treated mouse, whereas the IL- 15 mouse exhibited p24 levels comparable to those of the control (**Extended Data Fig. 11a**). These changes were accompanied by decreased CD4⁺T-cell levels and increased CD8⁺T-cell levels (**Extended Data Fig. 11a**).

The overall analyses of the frequency of p24^+^CD4^+^T cells including all mice with enough cells in each arm to be considered for comparative analyses showed no significant differences in spleen or blood (**Fig. 7d**). Of note, the individual analysis of the frequency of p24^+^ T cell in the spleen across the different PWH donors, revealed a broad inter-patient variability in the level of HIV infection remaining 2 weeks after administration of reactivated CD4^+^T cells, yet consistently showed reduced p24^+^CD4^+^T cell levels in the sequentially-treated group compared to controls (**Extended Data Fig. 11b**). Remarkably, intra-patient paired comparison showed that the frequency of reactivated p24^+^CD4^-^T cells in spleen was significantly lower in the sequential treatment group compared to the IL- 15 group (p=0.0313; **Fig. 7d**). Immunofluorescence analyses of CD4^+^p24^+^ cells in spleen confirmed results observed for the #PB02-derived samples (**Fig. 7e**), however, the low number of human cells detected in other samples limited detection of p24 in the rest of the samples (**Fig. 7f**). In addition, the proportion of CD8⁺CD3⁺ double-positive cells relative to the tissue area was highly variable but generally low, reaching a maximum of 30% in two IL-15/TGF-β1-treated recipients and 10% in three recipients from the IL-15 group (**Fig. 7f**, **right**). These results were supported by the quantification of total proviral HIV-DNA levels in spleen, which confirmed lower median levels in the IL-15/TGF-β1 arm, compared to the other groups (**Extended Data Fig. 11c**). Of note, HIV-DNA splenic detection was overall challenging due to low CD4^+^T cell counts, yet HIV-DNA was detectable in 4 out of 9 mice of the IL-15 group (44%) compared to 2 out of 7 (29%) from the sequential treatment group (**Extended Data Fig. 11c**). Regarding donor #PB02, viral DNA copies in the spleen of the IL-15/TGF-β1 mouse was 10^3^ lower than in their corresponding controls, which showed levels up to 10^7^ HIV-DNA copies per million of CD4^+^T cells (circle, **Extended Data Fig. 11c**).

Last, we evaluated the expression of some of the molecules of interest in CD8^+^T cells (PD-1, KLRG1, TIGIT; 4-1BB, CD69, CD103, CD39; GzmB; **Fig. 7g and Extended Data Figs. 11d-g**), which showed similar expression between IL-15 and sequential treatment groups in spleen and blood samples, except for a lower expression of TIGIT in sequentially treated CD8⁺T cells compared to IL-15 in blood (**Fig. 7g**), while 4-1BB and CD39 tended to be higher in samples from the IL-15 group compared to the sequential group (**Extended Data Figs. 11d and e**). In addition, in most animals, blood and splenic CD8^+^T cells showed higher frequencies of GzmB in sequentially-treated samples compared to IL-15-only treated samples from the same patient (**Fig. 7g and Extended Data Fig. 11f**). Of note, most murine samples derived from both CD8- conditions showed a significant enrichment of GzmB^+^ cells in the CCR7⁻CD8⁺T fraction (**Extended Data 11g-h**). Last, the frequencies of p24⁺CD4⁺ T cells across both CD8- groups significantly correlated with CCR7⁻CD8⁺T cells and 4-1BB⁺CCR7⁻CD8⁺ T cells (**Fig. 7h**). In summary, in successfully engrafted humanized mice, autologous IL- 15/TGF-β1-treated CD8⁺T cells exhibited superior control of reactivated HIV⁺ CD4⁺T cells from PWH compared to IL-15-only treatment.

## Discussion

Enhancing the antiviral capacity of CD8^+^T cell responses within tissue compartments supporting viral persistence at the time of ART discontinuation will be necessary to effectively limit the nascent viral rebound^3–5^. Here, we reveal that treating PBMC from PWH *ex vivo* with signals associated with residency establishment^7,8^ induces proliferation and remodeling of effector CD8^+^T cell subsets characterized by increased CD103 and CD39 expression, which identify effector cells that are more efficient at eliminating the reactivated reservoir, including intact viruses. Among the multiple benefits identified by sequential IL-15/TGF-β1 treatment, enhanced clonotype diversity and improved mitochondrial fitness of less terminal/exhausted effector phenotypes, may be recognized as a major advantage. Concordantly, adoptive transfer of these cytokine-treated CD8^+^T cells in PWH, as we preliminary tested in a humanized mouse model, holds promise as an immunotherapy to promote immune control and infection remission after treatment interruption.

The effect of treating T cells with IL-15 and/or TGF-β1 has been explored in multiple models underscoring the complexity of the resulting effect based on the cell subtype and model used, or the time of the intervention after infection^42,48–55^. IL-15 is essential for maintaining survival and homeostasis of memory CD8^+^T cells^49^. Spontaneous HIV control in memory CD8^+^T cells from non-controllers can be metabolically reprogrammed through IL-15 treatment *in vitro*^33^. IL-15 promotes fatty acid uptake, decreases glucose dependency and promotes mitochondrial respiratory capacity of bulk CD8^+^T cells, ultimately increasing suppressive capacity of HIV-specific CD8^+^T cell responses^33^. Treating PBMC with IL-15 induces cycling GzmB^+^CD8^+^T cells^54^, yet in untreated PWH it drives bystander activation of CD8^+^T cells, which predicts disease progression^54^. The observation that some mouse recipients presented higher levels of HIV infection in spleen in the IL-15 group could suggest this bystander activation, also reinforced by the high levels of CD69 expression observed in samples from PWH in IL- 15-treated samples, without an increase in the specific response. Moreover, out of the effector population, IL-15 only treatment, appeared to expand more exhausted signatures^21,22,40^ compared to sequential treatment, as detected by our scRNAseq analyses and confirmed by high expression of TIGIT in samples from PWH -including the ones from the mouse model- along with ROS accumulation. In this sense, the metabolic features of T cell exhaustion during chronic infection point to mitochondrial function as crucial for maintaining T cell function during persistent antigenic stimulation ^46, 47, 52^.

Here we show that combining IL-15 stimulation with sequential exposure to TGF- β1 further enhances HIV control, also linked to metabolic reprogramming and mitochondrial function. Sequential treatment induced epigenetic remodeling of the predominant expanded cluster, characterized by expression of *ITGA1* and *ITGAE* together with selective expression of *ENTPD1* and *TGFBR2,* the latter being a key regulator of TRM persistence and mitochondrial fitness^43^. Concordantly, after sequential treatment, exhausted CD39^+^ cells displayed reduced mitochondrial stress along with increased Ag- specific response. In addition, IL-15/TGF-β1-expanded clusters showed increased expression of the mTORC2 associated gene *RICTOR*, a gene related to the mTORC2 pathway, involved in cell survival and cytoskeletal organization^33^. This pathway, together with other genes enhanced by our sequential treatment (*BCL2, PRKCA, EOMES*), have been associated to natural control of HIV^33^. Other murine lymphocytic choriomeningitis virus models have also shown the involvement of TGF-β signaling repressing mTOR signaling in exhausted T cells and modulating the metabolism and function of exhausted precursors^51^.

In addition, TGF-β1 inhibits activation-induced cell death to allow the clonal expansion of effector T cells and the generation of memory T cells during immune responses^56^. Our data support the clonal expansion of effector CD8⁺T cells, accompanied by the induction of less differentiated intermediate memory effector phenotypes that have previously been shown to exhibit superior efficacy following adoptive transfer^57^. In our study, these cells were characterized by upregulation of CD27 and IL7R together with low KLRG1 expression. Prolonged ART has been linked to the rejuvenation of dominant CD8⁺T cell populations, characterized by the replacement of clonotypes displaying signatures of exhaustion and terminal differentiation with newly dominant clonotypes enriched for stem-like and early differentiation programs, features associated with natural control of viral replication^58^. In addition, control of viral rebound after an intervention in PWH is associated with superior pre-intervention HIV-specific CD8^+^T cell proliferative capacity and signatures of metabolic fitness and reduced T cell exhaustion ^59^, supporting the potential impact of our immunotherapy.

Expression of immune checkpoints in Ag-specific CD8^+^T cells in settings of chronic antigen stimulation may be characteristic of exhausted profiles, which can also infiltrate and reside in tissues, acquiring transcriptional profiles partially shared with TRM^18,21–23^. However, bona fide TRM cells, even under homeostatic conditions, also share key features with dysfunctional exhausted T cells, including immune checkpoint molecules^8,9,18,38^. Indeed, a recent report indicates that PD-1 signaling mediates skin CD8^+^TRM cell specification, lineage fate commitment, tissue niche engraftment and TGF- β responsivity^60^. Still, in our study, we can only relate expanded subsets to phenotypes similar to ‘bona fide TRM cells’ or ‘TRM-like cells’^18^, since neither migration to peripheral tissues nor residence was assessed. In this sense, the low frequencies of CD8^+^T cells detected in our mouse model, particularly of the CCR7^-^ fraction, in IL-15/TGF-β1-treated CD8⁺T-cell recipients could actually indicate, among other explanations, non-lymphoid tissue infiltration. However, no other tissues beyond spleen were assessed. Future studies will inform on the capacity to infiltrate to which tissues and for how long these cells persist after immunotherapy. Nevertheless, in the setting of a programmed ATI, if ART withdrawal and adoptive cell transfer are optimally timed, prolonged persistence of these cells within tissues may be unnecessary. Local inflammatory signals arising from early viremic foci would be expected to recruit administered effector CD8⁺T cells. Consistent with this model, we found that these cells were enriched for CXCR3 expression, supporting the potential for a chemokine-mediated recruitment ("pull") effect^61^.

We acknowledge that our study has several limitations. First, we performed scRNAseq in multiple samples from a single donor. Large hysterectomy samples from HIV women are not common in our settings, which together with the cost of RNAseq analyses limited including multiple donors. However, we were able to confirm multiple findings from gene expression data, both in blood conditions and in cervical tissue, validating scRNAseq data. Another limitation of this study was the animal model, which is subject to substantial variability in engraftment efficiency and therefore requires large cell numbers and multiple recipient replicates to validate the findings. While the number of cells administered was on the low end, and we detected high variability, a consistent benefit of the sequential treatment over the other conditions was detected. Perhaps administering both CD4^+^ and CD8^+^T cells by another route^62^ would improve engraftment. Last, a major limitation is that we did not assess other tissues beyond blood and spleen in this model, restraining from detecting viral containment in non-lymphoid tissues were most likely sequentially-treated cells performed.

Overall, we identify a strategy that improves the antiviral CD8^+^T cell response against persisting Ag by expanding reinvigorated cells with enhanced tissue-access and residency potential. This strategy can certainly be combined with optimal epitope stimulation approaches ^59^ and, in certain patients, with the blockade of molecules that represent breaks of an effective response (i.e. PD-1). Of note, such interventions need to be well timed and contextualized in order to avoid harmful auto-aggressive T cell responses^60^. Still, some of the CD8^+^T cell qualities modified by this treatment align with desired features for HIV control highlighted in recent studies^58,59^. Ultimately, we propose the induction of clonally diverse and reprogrammed CD8^+^TRM-like phenotypes expressing CD103 and or CD39 by sequential IL-15/TGF-β1 treatment, as a promising CD8^+^T cell-based immunotherapeutic strategy to enhance control of rebound viruses and elimination of persistent antigens like HIV-1.

## Methods

### Study subjects and samples

Blood and cervical tissue samples from ART-treated HIV^+^ women were obtained from hysterectomies or cone biopsies at the Hospital Universitari Vall d’Hebron (HUVH), as part of cervical dysplasia and cancer screening preventive treatment. Additional blood from HIV-1 infected men and women under ART attending the HIV unit of the HUVH processed and stored in a collection (National Biobank register #C.0003590) were used. Fresh large blood donations from selected PWH were provided by the members of our HIV unit in collaboration with the Catalan Blood Bank. **Extended Data Table 1** (*Not available for a preprint*) displays PWH information regarding sex, age, time since HIV diagnosis, plasma viral loads, CD4^+^T cell counts, time of ART-suppression and treatment. All the participants in this study signed a written informed consent before sample collection and study protocols were approved by the corresponding Ethical Committee (Institutional Review Board numbers PR(AG)270/2015, PR (IR) 294/2017 and PR(AG)192/2023).

### Antibodies

The following directly conjugated antibodies from BD Biosciences, unless otherwise stated, were used in flow cytometry experiments: **Cervical phenotype panels:** anti-HLA-DR-PerCP-Cy5 (G46-6), anti-PD-1-BV421 (EH12.1anti-CD69-Horizon-PE- CF594 (FN50), anti-CD4-BV605 (RPA-T4), anti-CD19-V500 (HIB19), anti-CCR7-PE (3D12), anti-CD49d(α4)-FiTC (9F10), anti-β7-APC (FIB504), anti-CD39-APC-Cy7 (A1, eBiosciences), anti-CD3-eVolve 655 (OKT3, eBiosciences), anti-CD363(S1PR1)- eFluor 660 (SW4GYPP, eBiosciences), anti-CD45-Alexa Fluor 700 (Hl30, BioLegend), anti-CD103-FiTC (BER-ACT8, BioLegend), anti-CD49a(α1)-PE (TS2/7, BioLegend), anti-CXCR3-FiTC (G025H7, BioLegend), anti- IL-7R-FiTC (A019D5, BioLegend), anti-CD132-PE (VI C-89, BioLegend), anti-CD122-APC (TU27, BioLegend), anti- CCR2-PE (48607, R&D Systems), anti-TCR-γδ-FiTC (11F2, Miltenyi Biotec), anti- CD161-PE (191B8, Miltenyi Biotec), anti-T-bet-BV421 (4B10, BioLegend), anti- Eomes-PE-Cy7 (WD1928, eBioscience) and anti-Hobit-AF647 (Sanquin-Hobit/1); **Intracellular cytokine staining**: anti-CD107a-PE-Cy5 (H4A3), anti-PD-1-BV421 (EH12.1), anti-CD69-PE-CF594 (FN50), anti-CD8-BV650 (RPA-T8), anti-CD45- BV605 (HI30), anti-CD3-PE-Cy7 (SK7), anti-CD39-APC-Cy7 (A1, BioLegend), anti- CD103-FITC (BER-ACT8, BioLegend), anti-IFNγ-AF700 (B27, Invitrogen), anti- perforin-PE (B-D48, Biolegend); **Blood and cervical tissue from HIV^+^ women sorting for functional assay:** anti-PD-1-BV421 (EH12.1), anti-CD69-PE-CF594 (FN50), anti- CD19-V500 (HIB19), anti-CD4-BV605 (RPA-T4), anti-CCR7-PE (3D12), anti-CD8-APC (RPA-T8), anti-CD3-PerCP (SK7), anti-CD32-PE-Cy7 (FUN-2, BioLegend), anti- CD39-APC-Cy7 (A1, eBiosciences), anti-CD45-Alexa Fluor 700 (Hl30, BioLegend), CD103-FiTC (BER-ACT8, BioLegend); **Phenotyping of expanded PBMCs:** anti- CCR7-PE (3D12), anti-CD3-PerCP (SK7), anti-CD4-BV605 (RPA-T4), CD8-APC (RPA-T8) anti-CD69-PE-CF594 (FN50), anti-CD39-APC-Cy7 (A1, BioLegend), anti-CD103-FiTC (BER-ACT8, BioLegend), anti-EOMES-PE-Cy7 (WD1928, Invitrogen), anti-Ki-67-AF700 (SolA15, Invitrogen), anti-S1PR1-eF660 (SW4GYPP, Invitrogen), and anti-T-bet-BV421 (4B10, BioLegend), anti-CXCR3-BV650 (G025H7 BioLegend) and anti-CD49a(α1)-PE (TS2/7, BioLegend); **p24 expression in co-cultures:** anti-CD3- PerCP (SK7), anti-CD8-FITC (HIT8a), anti-p24-PE (FH190-1-1, Beckman Coulter). **Sorting of CD8^+^ and CCR7^-^CD8^+^ subpopulations for IPDA and for scRNAseq:** anti- CCR7-PE (3D12), anti-CD3-BV650 (UCHT1), anti-CD4-BV605 (RPA-T4), CD8-APC (RPA-T8), anti-CD69-PE-CF594 (FN50), anti-CD39-APC-Cy7 (A1, BioLegend), anti-

CD103-FiTC (BER-ACT8, BioLegend), anti-CD45-Alexa Fluor 700 (Hl30, BioLegend), anti-PD-1-BV421 (EH12.1), anti-CD137-PE-Cy7 (4B4-1, BioLegend). **Verification panel cervical sample and PBMCs:** anti-CD45-BV605 (HI30), anti-CD3-PE-Cy5 (UCHT1, BioLegend), anti-CD8-APC (RPA-T8), anti-CCR7-AF647 (150513), anti-IL7- Rα-FITC (A019D5, BioLegend), anti-CD69-PE-CF594 (FN50), anti-CD103-BV650 (BER-ACT8), anti-CD39-APC-Cy7 (A1, BioLegend), anti-CD137-PerCP-Cy5.5 (4B4-1, BioLegend), anti-CD49a-PE-Cy7 (TS2/7, BioLegend), anti-CD27-PE-eFluor610 (LG.7F9, Life Technologies), anti-CD52-SB702 (CF1D12, Life Technologies), anti- KLRG1-BV480 (13F12F2, Life Technologies), anti-TCR-γδ-PerCP-eFluor710 (B1.1, Life Technologies), anti-PD-1-BV421 (EH12.1), anti-LAG3-APC-Fire810 (11C3C65, BioLegend), anti-TIM3-SB780 (F38-2E2, Life Technologies), anti-TIGIT-eFluor450 (MBSA43, Life Technologies), anti-CTLA-4-RB780 (BNI3), anti-IFNγ-AF700 (B27, Invitrogen), anti-Granzyme-B-PE (GB11, Life Technologies). **Flow Cytometry analysis of mitochondrial ROS:** anti-PD-1-PercP (EH12.2H7, BioLegend), anti-CD8-PeCy7 (RPA-T8), anti-CCR7-APC (150503), anti-CD39-APC-Cy7 (A1, BioLegend), anti-

CD69-Pacific Blue (FN50), anti-CD103-BV510 (Ber-ACT8). **Sorting of CD8^+^ T cells from PBMC of PWH for adoptive cell transfer to mouse recipients (batch 2):** Anti- CD45-AF700 (Hl30, BioLegend), anti-CD3-BV650 (UCHT1), anti-CD4-BV605 (RPA- T4), anti-CD8-APC (RPA-T8), anti-CD56-BV711 (NCAM16.2), anti-CCR7-PE (3D12), anti-CD69-PE-CF594 (FN50), anti-CD103-FITC (BER-ACT8, BioLegend), anti-CD39-

APC-Cy7 (A1, BioLegend), anti-CD137(4-1BB)-PE-Cy7 (4B4-1, BioLegend) and anti-PD-1-BV421 (EH12.1). **Spectral flow cytometry analysis of blood and spleen of a humanized mouse model:** For batch 1: Anti-CD45-BV605 (HI30), anti-CD3-PerCP (SK7), anti-CD4-APC (OKT4, BioLegend), anti-CD8-BV650 (RPA-T8), anti-CCR7- BV711 (3D12, Thermo Fisher), anti-CD69-PE-CF594 (FN50), anti-CD103-FITC (BER-

ACT8, BioLegend), anti-CD39-APC-Cy7 (A1, BioLegend), anti-4-1BB-PE-Cy7 (4B4- 1, BioLegend), anti-PD-1-BV421 (EH12.1), anti-IFNγ-AF700 (B27, Invitrogen), anti-Granzyme-B-AF647 (GB11) and anti-p24-PE (FH190-1-1, Beckman Coulter). For batch 2: the same antibodies were used despite the change of anti-CCR7-AF700 (150513), the exclusion of anti-IFNγ-AF700 (B27, Invitrogen) and the addition of anti-KLRG1-APC- Fire810 (14C2A07, BioLegend) and anti-TIGIT-PerCP-eFluor-710 (MBSA43, Life Technologies). **IF analysis of spleen tissue staining:** Anti-CD8-Opal Fluorophore 570 (CAL67, Abcam), anti-CD3-Opal Fluorophore 520 (CAL54, Abcam) and anti-p24-Opal Fluorophore 650 (5, Abcam).

### Cervical tissue digestion and flow cytometry phenotyping

As previously described by our group^23^, endocervical and ectocervical samples were obtained in antibiotics-containing RPMI 1640 medium once the Pathology Service had confirmed the healthy status of the tissue. Within the 24h following surgery, the mucosal epithelium and the underlying stroma were isolated from the muscular tissue and dissected into approximately 2-mm^3^ blocks. Blocks were enzymatically digested with 5 mg/ml collagenase IV (Gibco) and the resulting cellular suspension was washed twice and stained for viability with Live/Dead Aqua (Invitrogen). After washing, cells were separated in different tubes and stained as described above (see “Antibodies” section). Stability after collagenase IV treatment was ensured for each cell surface marker on PBMCs as described ^23^.

### PBMC cytokine treatment

Phenotype and function of PBMC from PWH under suppressive ART were analyzed in different conditions, depending on the assay: 1) baseline; 2) 6-day culture without stimulus; 2) 3-day culture with 50 ng/mL of IL-15 (Miltenyi Biotec.); 4) 3-day culture with 50 ng/mL of IL-15 and 1µg/mL of clade B Gag peptides; 5) 3-day culture with 50 ng/mL of IL-15 and 3-day culture with 50 ng/mL of TGF-β1 (R&D); 6) 3-day culture with 50 ng/mL of IL-15 and 1µg/mL of clade B Gag peptides and 3-day culture with 50 ng/mL of TGF-β1 (R&D). For all conditions, cells were cultured with R10 supplemented with 10 U/ml of IL-2.

### Detection of Gag-specific CD8^+^T cell response

Cervical biopsies and blood from HIV^+^ people under ART were obtained. Tissue samples were digested following the same protocol described above and PBMC were obtained from blood. Individual or matching samples from WWH were stimulated with 5 µg/mL of clade B Gag peptides (NIH AIDS Reagent Program), DMSO (negative control, Fisher scientific) or 1 μM ionomycin and 10 ng/mL PMA as a positive control. In some cases, 10 µg/ml Nivolumab (Vall d’Hebron Pharmacy Department) or 100 µM ARL67156 (Sigma-Aldrich; A265) were added during Gag-stimulation to block PD-1 or CD39, respectively. Following standard intracellular cytokine staining protocols, cells were stimulated for 5.5 h at 37°C with R20 in the presence of 3 μL/mL α-CD28/CD49d (clones L293 and L25), 1 μL/mL Brefeldin A, 0.7 μL/mL Monensin and 50 μL/mL anti-CD107a-PE-Cy5 (all from BD Biosciences), since its expression is very rapid. Cellular suspensions were stained for Live/Dead Aqua (Invitrogen) and as described above for phenotype and intracellular cytokine staining (see “Antibodies” section). Staining of Eomes, Ki-67, S1PR1 and T-bet was performed by using Foxp3/Transcription Factor Staining Buffer Set from Invitrogen according to manufacturer’s protocol. After fixation in PBS 2% PFA, cells were acquired in a BD LSRFortessa^TM^ flow cytometer (VHIR) and analyzed with FlowJo vX.6.1 software.

### Determination of p24 after viral reservoir reactivation

Cryopreserved PBMC from PWH under suppressive ART were cultured for 6 days in R10 with 10 U/mL of IL-2 for the control condition, with sequential stimulation with 50 ng/mL of IL-15 (3 days) and 50 ng/mL of TGF-β1 (next 3 days) for the IL-15/TGF-β1 sequential treatment condition, or only for the 3 first days for the IL-15 condition, as described in the previous section. The day before co-culture set up, PBMC from the same patient were thawed to isolate CD4^+^T cells using MagniSort Human CD4 T Cell Enrichment Kit (Invitrogen) and were incubated with R10 supplemented with 1 µM Raltegravir, 1 µM Darunavir, 1 µM Nevirapine and 10 µM Q-VD-Oph (Selleckchem) for 2 h. Then, ¼ CD4^+^T cells were separate to use as a reactivation control and the remaining cells were reactivated using 100nM Ingenol for 22 h as a LRA. Next day, CD8^+^T cells were isolated from different expanded PBMC conditions using CD8^+^T Cell Isolation Kit (Miltenyi Biotec) and co-cultures (1:1) were established in a 96-well plate at 1.5 × 10^6^ cells/mL: 1) ∼10^6^ CD4^+^ LRA^-^, 2) ∼ 500,000 CD4^+^ LRA^+^, 3) ∼500,000 CD4^+^ LRA^+^ with ∼500,000 CD8^+^, 4) ∼500,000 CD4^+^ LRA^+^ with ∼500,000 CD8^+^ from IL-15-expanded culture and 5) ∼500,000 CD4^+^ LRA^+^ with ∼500,000 CD8^+^ from sequential IL-15/TGF-β1 expanded culture (exact cell numbers may have differed inter-patient but not intra-patient). After 4h, cells were stained for viability with Live/Dead Fixable Far Red Dead Cell Stain Kit (Invitrogen) and for p24 expression panel (see “Antibodies“ section). After fixation in PBS 2% PFA, cells were acquired in a BD FACSCalibur^TM^ flow cytometer (VHIR). The limit of detection of this analysis was 9.6 molecules/10^6^ CD4^+^T cells, which corresponded to three times the standard deviation of the number detected in uninfected individuals. This threshold was established by measuring p24 expression in fifteen uninfected individuals before and after *in vitro* infection.

### Functional assay to determine Intact proviral DNA assay

Cryopreserved PBMC from PWH under suppressive ART were thawed to isolate CD4^+^T cells using Dynabeads CD4 Positive Isolation Kit (Invitrogen) and were *ex vivo* infected with HIV-1 BaL by spinoculation and cultured in R10 supplemented with 100U/ml IL-2 (Merck), between 1- 2 × 10^6^ cells were not infected for the negative control. In the first functional assay, the isolated CD4 fraction was cryopreserved after isolation with subsequent *ex vivo* infection with HIV-1 BaL as specified before or cryopreserved after *ex vivo* superinfection. The CD4^-^ fraction was cultured for 3 days in R10 supplemented with 10 U/mL of IL-2 and IL-15 and for 6 days in R10 supplemented with 10 U/mL of IL-2 and sequential stimulation with IL-15 and TGF-β1 to expand tissue resident-associated phenotypes as described before. On day 3 and 6, cells were cryopreserved. One day before cell co-culture, the CD4 fraction and both CD4^-^ fractions, IL-15 or with sequential stimulation, were thawed and rested overnight in R10 with 10 U/mL of IL-2. Then, the stimulated CD4^-^ T cells were stained for viability with Live/Dead Aqua (Invitrogen) and surface markers as described above (see “Antibodies” section). After staining, CCR7^-^ and CCR7^+^ subpopulations from live CD45^+^CD3^+^CD8^+^ cells were sorted in a sorter Cytek Aurora 5L spectral cytometer (VHIR) from each condition (IL-15 and sequential stimulation).

For the second functional assays addressing specific effector CD8^+^T cell subsets after sequential treatment, the CD4^-^ fraction was cultured for 6 days in R10 supplemented with 10 U/mL of IL-2 and sequential stimulation with IL-15 and TGF-β1 to expand tissue resident-associated phenotypes as described before. On day 6, these cells were stained for viability with Live/Dead Aqua (Invitrogen) and surface markers as described above (see “Antibodies” section). After staining, four subpopulations of Live CD45^+^CD3^+^CD8^+^CCR7^-^ cells were sorted in a sorter Cytek Aurora 5L spectral cytometer (VHIR): 1) CD69^+^ CD103^+^ (conventional TRM-like), 2) CD69^+^CD103^-^ CD39^+^(unconventional TRM-like), 3) CD69^+^CD103^-^CD39^-^ and 4) CD69^-^CD103^-^CD39^-^. Then, in both functional assays, CD4^+^T HIV-infected cells were co-cultured with each CD8^+^ sorted subpopulation in a 96-well plate at 1 × 10^6^ cells/mL (R10) for 4h. Target to effector ration was between 1:1 and 1:6 based on patient, always maintaining the same ratio between intra-patient conditions. Baseline CD4^+^T cells and CD4^+^T cells infected *ex vivo* were used as controls. After 4h of incubation at 37°C in 5% CO2, the remaining viable CD4^+^ T cells were isolated from each condition using a combination of the EasySep™ Dead Cell Removal (Annexin V) Kit (StemCell Technologies) and the MagniSort Human CD4 T Cell Enrichment Kit (Invitrogen) and pellets from each condition were obtained and freezed at -80°C. Cell pellets were then lysed with proteinase K-containing lysis buffer and incubated at 55°C overnight, followed by a 95 °C incubation for 5 min, to obtain DNA for HIV-1 Intact Proviral DNA Assay (IPDA) described below.

### Intact proviral DNA assay (IPDA)

was performed as described before and using primers and probes specific for the Ψ HIV gene (HIV-1 Ψ forward 5′- CAGGACTCGGCTTGCTGAAG-3′, HIV-1 Ψ reverse GCACCCATCTCTCTCCTTCTAGC and probe 5′ 6-FAM-TTTTGGCGTACTCACCAGT-MGBNFQ-3′) and env HIV gene (HIV-1 env forward 5′-AGTGGTGCAGAGAGAAAAAAGAGC-3′, HIV-1 env reverse 5′-GTCTGGCCTGTACCGTCAGC-3′, HIV-1 env intact probe 5′-VIC- CCTTGGGTTCTTGGGA-MGBNFQ-3′, and HIV-1 anti-Hypermutant env probe 5′- CCTTAGGTTCTTAGGAGC-MGBNFQ-3′). The hRPP30 gene was used for cell input normalization and to quantify DNA shearing. Samples were analyzed in a QIAcuity One 2-plex System (Qiagen).

### Single cell RNA sequencing sample processing

PBMC from a single donor HIV^+^ ART- suppressed subjected to a hysterectomy (#M47) were directly stimulated overnight in 96 round well-plate with 5ug/ml clade B Gag peptides or DMSO (negative control, Fisher scientific) in R10 with 3uL/mL FastImmune α-CD28/CD49d (clones L293 and L25, BD Biosciences) at 7 x 10^6^ cells/mL for baseline condition, after three days stimulation with the IL-15 together with1ug/ml clade B Gag peptides in R10 10U/mL IL-2 (Merck) at 1.5 x 10^6^ cells/mL for the IL-15 only condition, or after sequential IL-15+Gag followed by 3-day culture with TGF-β1 as described before. The next day, each condition was stained for viability with Live/Dead Aqua (Invitrogen) and surface markers as described above (see “Antibodies” section). For the PBMC samples, live CD45^+^CD3^+^CD8^+^CCR7^-^ cells were sorted by using Cytek Aurora 5L spectral cytometry sorter (VHIR).

Additionally, cervical tissue from the same donor was digested and stimulated with Gag overnight as previously described for PBMC samples. Next day, cervical cell suspension was stained with the same panel and total live CD45^+^ cells were sorted following the same protocol as for PBMC. A cervical sample from the first uninfected donor available who underwent hysterectomy was used as control by overnight stimulation with DMSO before sorting as performed for the HIV^+^ donor. All PBMC and cervical samples were immediately sorted in low retention tubes containing 4°C PBS + BSA 0.04% and subsequently centrifuged at 600g for 8min at 4°C to suspend again to desired concentration (700-1200 cells/µl) on the same buffer, before transport on ice to the Single Cell Unit of the Josep Carreras Leukaemia Research Institute (IJC).

### Library preparation and sequencing

Single cell suspension was immediately loaded in the Chromium Next GEM Chip G wells, to have a final recovery of approximately 10,000 cells per suspension. The Chromium Controller performed single cell partitioning and barcoding. Cells were partitioned into nanoliter-scale Gel Beads-in-emulsion (GEMs) where all generated cDNA shared a common 10x Barcode. A pool of ∼3,500,000 10x Barcodes were sampled separately to index each cell’s transcriptome. Dual Indexed libraries were generated and sequenced from the cDNA and 10x Barcodes were used to associate individual reads back to the individual partitions. Single Cell 3′ v2 libraries were generated following the Single Cell 3′ v2 Reagent Kits User Guide (Document CG00052, 10X genomics) and libraries obtained sequenced on the Illumina NovaSeq X platform at 40,000 reads/cell.

### scRNA-seq analysis

These analyses were performed by the Statistics and Bioinformatics Unit (UEB) at VHIR. Sequencing reads were checked for optimal quality using FASTQC tool v0.11.9 and aligned to human reference genome GRCh38 with Cell Ranger v7.2.0. The cellranger multi pipeline was used to jointly process GEX and VDJ libraries for each sample, which performs alignment, filtering, barcode counting, and UMI counting on the Gene Expression libraries. It also performs sequence assembly and paired clonotype calling on the V(D)J libraries. For cervical samples, because no clear steep cliff was observed in the Barcode Rank Plot obtained with Cell Ranger for sample from #M47, DropletQC R package (Muskovic et al 2021) was used to filter out empty droplets. Downstream analyses were mainly performed in the statistical language R v4.2.0 (Bioconductor v3.15). After alignment and quantification of reads to annotated mRNAs, we applied different quality checks to detect low quality cells. Based on quality control results, mild filtering was performed on the number of features per cell and mitochondrial/ribosomal counts. A threshold of < 20% mitochondrial counts, > 5% ribosomal reads, > 250 features and > 30,000 UMI counts per cell was used. Datasets were normalized using SCTransform algorithm within Seurat v4.3 ecosystem and integrated using Harmony algorithm (Korsunsky et al 2019) based on the first 30 PCs. Cluster analysis was performed using the Louvain clustering algorithm at resolution of 0.5 and visualized on top of UMAP embeddings. Cell type markers were identified for each cluster using FIndAllMarkers function from Seurat with default parameters (Wilcoxon rank sum test) and used to annotate clusters manually. Differences in gene expression across conditions were assessed with FindMarkers function from Seurat (two-sided Wilcoxon rank sum test). Genes with a FDR-adjusted P value < 0.05 were considered significantly upregulated or downregulated if > or < 0.25-fold change (FC) respectively. Volcano plots were generated using the processed data and plotted using VolcaNOseR v1.0.3 (https://huygens.science.uva.nl/VolcaNoseR/). The analysis of biological significance was based on enrichment analysis over Gene Ontology database (GO, Biological Process category) using clusterProfiler R package. Additionally, developmental trajectories and pseudotime relationships were inferred using Slingshot v2.4 R package. TCRαβ repertoires in samples were analyzed using scRepertoire v2.0.0 R package (Borcherding et al., 2020). Clonal expansion was defined by cells sharing CDR3 nucleotide sequences, V gene, and J gene usage. Clonotypes were defined using the strict criterion and categorized according to their expansion as Single, Small, Medium, Large, and Hyperexpanded. Shannon index indicating repertoire diversity was used to estimate the richness of each sample and Jaccard similarity index implemented in the scRepertoire R package was used to quantify TCR clonal repertoire overlap between cell clusters.

### Flow Cytometry analysis of mitochondrial ROS

PBMC from ART-suppressed PWH were expanded or not as described in the ‘*Sample processing for Single cell RNA sequencing’* section and overnight exposed to 5 ug/ml clade B Gag peptides or DMSO (negative control) and cryopreserved. Frozen samples were shipped to Hospital Universitario La Princesa, Madrid, were 1.5 - 2 million cryopreserved PBMC were rested in complete RPMI 10% FBS media for 2 h after thawing. Afterwards, cells were simultaneously incubated with 5µM MitoSox and 25nM MitoTracker probes (Invitrogen) in a 1X PBS solution containing 0.5% BSA and 2mM EDTA for 15 mins at 37 °C. After washing the probes with an excess of the same buffer, samples were stained for surface markers as described above (see “Antibodies” section). After staining, samples were acquired using a BD FACSCanto™ II cytometer.

### Spectral flow cytometry validation analyses. Spectral flow cytometry validation analyses

PBMC from ART-suppressed PWH (n=8, **Extended Data Table 1**) at the three blood study conditions (Baseline, IL-15+Gag and IL-15+Gag/TGF-β1) and concomitant cervical cone biopsy from four of them were obtained. PBMCs were expanded or not as described in the ‘*Sample processing for Single cell RNA sequencing’* section and samples were cryopreserved at day 0, 3 and 6 for each condition, respectively. Cervical samples were digested as described above and cell suspensions were also cryopreserved. After thawing and recovery, all samples were stimulated overnight in 96 round well-plate with 5ug/ml clade B Gag peptides or DMSO, as described above. Next day, samples were stained for viability with Live/Dead Aqua (Invitrogen) and surface markers as described above (see “Antibodies” section). Intracellular staining of Granzyme B and IFNγ was performed by using the FIX and PERM Cell Permeabilization kit (ThermoFisher) according to manufacturer’s protocol. After fixation in PBS 2% PFA, samples were acquired in a Cytek Aurora 5L spectral cytometer (VHIR).

### Humanized mouse model

All mouse experiments were conducted in a Biosafety Level 3 (BSL3) facility at the Comparative Medicine and Bioimage Centre of Catalonia (CMCiB). Two different NOD scid gamma mice (NSG) mice variants were used for this animal mice model, NOD scid Il2rynull B2mnull mice (Batch 1; NSG B2m; strain 010638) and standard NSG mice (Batch 2; strain 614) were procured from the Jackson Laboratory. From a total of 44 male mice, 22 mice were procured for each batch. Animal studies were performed by trained researchers and approved and assessed by the Committee on the Ethics of Animal Experimentation from the CMCiB (CSB-23-003) with the authorization of the Generalitat de Catalunya (code: 12009). All procedures involving experimental animals were carried out in compliance with pertinent institutional and national animal welfare laws, guidelines and policies.

Between 3-10 mice/patient were tested based on cell numbers for adoptive transfer (CD4 and CD8^+^T cells). Based on this, mice were distributed among patient-derived groups in three conditions (**Extended Data Table 2**): 1. Control (without addition of CD8^+^ T cells), 2. IL-15 (with CD8^+^ T cells stimulated with IL-15) and 3. Sequential IL- 15/TGF-β1 (with CD8^+^ T cells stimulated with IL-15/TGF-β1). All mice were administrated with autologous reactivated CD4^+^ T cells together with a remaining fraction depleted from CD4, CD8 and NK cells.

Fresh large blood donations were processed as specified above to obtain PBMCs (see “Blood processing” section). Subsequently, CD4^+^T cells were isolated using the Dynabeads CD4 Positive Isolation Kit (Invitrogen), according to manufacturer’s protocol and cryopreserved. 3 days prior cell administration purified CD4^+^T cells were thawed and reactivated by CD3/CD28 stimulation. For that, U-96 well plates were previously coated with monoclonal anti-CD3 antibody (OKT3, Life technologies; at 5 ug/ml for batch 1 and 1.25 ug/ml for batch 2 in PBS) overnight and washed twice with PBS 1x before adding the CD4^+^ T cells, which were cultured overnight in these CD3-coated-plates with R10 and the monoclonal anti-CD28 antibody (CD28.2, Life technologies; at 5 ug/ml for batch 1 and 1.25 ug/ml for batch 2) at a cell concentration of 2.66 x 10^6^/ml. The non-CD4^+^T cell fraction was cultured and stimulated either with IL-15 or with sequential IL-15/TGF- β1 treatment as specified above (see “Cytokine treatment of PBMCs” section). After stimulation, cells were cryopreserved until the day before cell administration.

Batch 1 - CD8^+^ T cell isolation: On the day before cell administration and from both stimulated non-CD4^+^ T cell fractions, CD8^+^ T cells were isolated using the Dynabeads CD8 Positive Isolation Kit (Invitrogen) according to the manufacturer’s protocol. Then, purified CD8^+^ T cells from each condition (IL-15 and IL-15/TGF-β1) were cultured overnight in R10 and maintained at 37 °C in a 5% CO2 incubator. The non- CD4, non-CD8 T-cell fractions from each stimulated condition were independently processed using the CD56 Positive Selection EasySep™ Kit II (StemCell Technologies) to obtain a purified non-CD4, non-CD8, non-CD56 cell population (hereafter referred to as the remaining fraction).

Batch 2 - CD8^+^T cell isolation: For this batch, the only difference in the isolation of CD8 and the CD4/CD8/CD56-depleted is that the cells were obtained by spectral flow cytometry sorting. For that, the day before cell administration, both cytokine-stimulated non-CD4^+^T cell fractions were thawed and stained separately for viability and surface as specified in the “Flow cytometry phenotyping” and “Antibodies” sections. Once cells were properly stained, Aqua^-^ CD45^+^ CD3^+^ CD8^+^ T cells and Aqua^-^ CD45^+^ CD3^-^ CD56^-^ T cells (remaining fraction) from each condition (IL-15 and IL-15/TGF-β1) were sorted in a Cytek Aurora 5L spectral sorter (High Technology Unit, VHIR). All cells were immediately acquired in low retention tubes containing 4 °C PBS + BSA 0.04% and subsequently overnight culture in R10 and maintained at 37 °C in a 5% CO2 incubator. The remaining fraction from each condition were mixed together and cryopreserved until the day before adoptive transfer, when were thawed, cultured in R10 and maintained at 37 °C in a 5% CO2 incubator. The next day, the remaining fraction was mixed with autologous reactivated CD4^+^T cells for mouse administration.

Ratios between 1:1 and 2.2:1 of CD4:CD8 cells were maintained for each patient-derived mouse group (**Extended Data Table 2**) and resuspended in PBS for subsequent mouse administration (between 100 ul to 150 ul/mice). For the purified CD8^+^T fractions treated with IL-15 or sequential treatment, cells of each condition were resuspended in PBS for mouse administration (between 100 ul to 150 ul/mice).

### Humanized mouse model sample processing

Briefly, plasma and PBMCs were isolated from blood samples by centrifugation at 2000 g for 20 min at 4 °C. Plasma samples were then lysed with RAV1 buffer (NucleospinTM RNA Virus kit, Cultek) and preserved at - 80 °C for further analysis. Remaining red blood cells present in PBMCs were lysed using ACK Lysis buffer (Fisher Scientific) following manufacturer’s instructions. Spleens were excised and divided in three parts for flow cytometry, IF and total HIV-DNA determinations. Spleen tissue for flow cytometry and HIV-DNA analyses was minced into small pieces and processed through a 70 μm cell strainer (Falcon). The obtained PBMC and splenic cells were processed for spectral flow cytometry analysis by staining with viability, surface and intracellular markers as described in the “Antibodies” section. The spleen tissue excised for IF was fixed in PFA 4% for 4h at 4 °C and then, let in 70% ethanol at 4 °C until paraffin inclusion was performed as specified below (see “IF analysis of spleen tissue” section).

### Plasma Viral load (pVL) determination

Animal model batch 1: pVL was measured every three days. HIV RNA was extracted from plasma using Nucleospin^TM^ RNA Virus kit (Cultek, Macherey-Nagel) following the manufacturer’s instructions. Reverse transcription of RNA to cDNA was performed with SuperScript^TM^ III Kit (Invitrogen) in accordance with the instructions provided by the manufacturer, and cDNA was quantified by quantitative PCR (qPCR) using primers and probes specific for the 1-LTR HIV region (LTR forward 5′-TTAAGCCTCAATAAAGCTTGCC-3′ and LTR reverse 5′- GTTCGGGCGCCACTGCTAG-3′; LTR probe 5′- CCAGAGTCACACACCAGACGGGCA-3′), and TaqMan^TM^ Gene Expression Master Mix (Thermo Fisher). Samples were analysed in Applied Biosystems QuantStudio5 system. Quantification of DNA was performed using a standard curve, and values were normalized to 1 million CD4^+^T cells. Animal model batch 2: HIV pVL was also measured every three days to evaluate viral load in plasma samples from NSG mice in collaboration with the Microbiology Department of the University Hospital Vall d’Hebron. pVL was determined using the Kit Cobas HIV-1 5800/6800/8800 192T IVD (Roche Diagnostics) was used together with the Cobas system (limit of quantification 20 copies/mL).

### Phenotyping of blood and spleen samples from a humanized mouse model

Lysed blood and tissue samples were stained with a viability dye, then cells were washed with PBS 1X and cell surface stained. After that, cells were washed and intracellularly stained. Finally, cells were washed and fixed with 1% PFA in PBS. See for more information in the “Antibodies” section. Samples were acquired in a Cytek Aurora 5L spectral cytometer (High Technology Unit, VHIR). Data was analysed using OMIQ software from Dotmatics (www.omiq.ai, www.dotmatics.com).

### Quantification of cell-associated HIV-DNA by qPCR in splenic tissue

Dry pellets of splenic cells of each mouse were obtained and frozen at -80 °C. Importantly, as human CD4^+^T cells were not isolated before the HIV-DNA determination, values were normalized to frequency of CD4^+^T cells (%) out of total CD45^+^ Live cells determined by flow cytometry of each corresponding mouse spleen sample. All samples were immediately lysed with a proteinase K-containing lysis buffer (55°C over-night and afterwards 5 mins at 95 °C). Total HIV-DNA quantification was performed in cell lysates by qPCR using primers against HIV long terminal repeat (LTR forward 5’- TTAAGCCTCAATAAAGCTTGCC-3’ and LTR reverse 5’- GTTCGGGCGCCACTGCTAG-3’; LTR probe 5’- CCAGAGTCACACACCAGACGGGCA-3’). Total HIV-DNA was quantified using a standard curve and CCR5 gene was used for cell input normalization. The cycling parameters were 50 °C for 2 min, 95 °C for 10 min and then 95 °C for 15 s and 60 °C for 1 min for 55 cycles of amplification.

### Immunofluorescence analysis of spleen tissue

Histology tissue slides out of fixed spleen tissue samples were prepared by the ICTS-NANBIOSIS group (U20/FVPR, at VHIR). Formalin-fixed, paraffin-embedded tissue slides underwent overnight deparaffinization at 65 °C, followed by xylene and ethanol dilutions and fixation in 10% neutral buffered formalin. For p24, CD8 and CD3 staining (see more information in the “Antibodies” section), slides were placed in pH=9 antigen retrieval at 95 °C for 30 min. After cooling, slides were permeabilized for 10 min with TBST 1x 0.1% Triton 100x. Then, slides were blocked with 1× Block Opal buffer for 10 min, washed with TBS/Tween washing buffer and incubated with anti-p24 (1:250) antibody. Subsequently, 1× Opal Anti-Mouse + Rabbit HRP was applied and incubated for 45 min at room temperature using a 1:100 dilution of Opal Fluorophore 650 reagent in 1× Plus Manual Amplification Diluent, according to the manufacturer’s instructions (AKOYA Biosciences, NEL811001KT). p24 fluorophore stripping was performed for 30 min at 95 °C and pH 9, simultaneously working as the antigen retrieval step for CD8. Washes and incubations were performed as described before with a working dilution for anti-CD8 of 1:150and using Opal Fluorophore 570. After CD8 stripping (performed as for p24), slides underwent through the same procedure for CD3 staining (1:150), except for Opal Fluorophore 520. After CD3 stripping, slides were counterstained with DAPI (1:1,000) for 7 min, washed, and mounted with Fluoromount-G (Invitrogen, Thermo Fisher). Images were acquired at 20× original magnification using a wide-field multidimensional Thunder microscope (Leica, High Technology Unit, VHIR). Sample counts and frequencies were normalized by area (mm^2^). Image analysis was performed using ImageJ (NIH), where binary masks for each marker were predefined for final cell counting.

### Statistical analyses

Data were analyzed using Prism software (GraphPad v10.2.2). Non-parametric two-tailed Mann-Whitney rank tests, Wilcoxon matched-pairs signed rank test and one-way ANOVA with post hoc Dunn’s correction for multiple comparisons were used unless otherwise indicated. Correlation calculation between two parameters was performed using Spearman’s correlation test.

### Extended Data

Extended Data Table 1

Extended Data Table 2

Extended Data Fig. 1

Extended Data Fig. 2

Extended Data Fig. 3

Extended Data Fig. 4

Extended Data Fig. 5

Extended Data Fig. 6

Extended Data Fig. 7

Extended Data Fig. 8

Extended Data Fig. 9

Extended Data Fig. 10

Extended Data Fig. 11

## Acknowledgments

We would like to thank all the patients who participated in the study and their providers. The authors thank Marina Rierola from “Banc de Sang i Teixits” in Vall Hebron, and Drs. Mª Elena Suárez and Sabina Salicrú from the Gynecology department from HUVH for referral of patients and sample collection as well as Dr. M. Ferrer and Esther Camacho, from the Statistics and Bioinformatics Unit (UEB, at VHIR), Dr. C. Mata, Head of Single Cell Unit at Josep Carreras Leukaemia Research Institute (IJC) and Jorge Díaz and Rosa Ampudia from the Comparative Medicine and Bioimage Centre of Catalonia (CMCiB) for excellent technical assistance.

## Funding

This work was primarily supported by grants from the Spanish Health Institute Carlos III (ISCIII, PI20/00160), cofunded by ERDF/ESF, “A way to make Europe”/"Investing in your future", and a Gilead fellowship GLD21-00049 and GLD23- 00012 to M.G.. This work was additionally supported with grants from MCIN/AEI/10.13039/501100011033 and the European Union «Next Generation EU»/PRTR (CNS2022-135549 and PID2023-149204OB-I00) and the Fundació La Marató TV3 (201814 and 202112). C.M-P. was supported by a Ph.D. fellowship from the Agència Gestió Ajuts Universitaris i de Recerca (AGAUR/FI/2023). The funders had no role in study design, data collection and analysis, the decision to publish, or preparation of the manuscript.

## Authorship Contributions

M.G. conceived and supervised the project. C.M-P., I.T. A.B-M., J.G-E., E. M-G., M.J.B. and M.G. designed and carried out experiments. C.M-P., I.T. A.B-M., J.G-E., E. M-G., M.J.B. and M.G. analyzed and interpreted data. J.C., L.M-B., C.C-M., B.P., P.N., J.N., J.B., A.C., and V.F. selected study subjects and provided samples. C.M-P. and M.G. wrote the manuscript. All authors reviewed the manuscript.

## Competing interests

The authors declare no conflict of interest.

## Notes

### Competing Interest Statement

The authors have declared no competing interest.

### Summary of Updates

This updated version includes substantial new data and a refocusing of the study on cytokine-mediated CD8+ T-cell reprogramming for HIV control. The manuscript excludes some clinical analyses to avoid overinterpretation of data derived from limited tissue samples, while incorporates an expanded cohort to corroborate finding from scRNAseq analyses. It also includes new proof-of-concept studies in a humanized mouse model. The results provide further evidence that sequential IL-15 and TGF-beta1 stimulation enhances antiviral CD8+ T-cell properties and supports improved control of HIV rebound.

