## Supplementary Information for "Sequential IL-15 and TGF-β1 stimulation expands reprogrammed CD103⁺CD39⁺ effector CD8⁺T cells and enhances HIV control"

Mancebo-Pérez et al.

**Extended Data Table 1.** Clinical data of people living with HIV included in the study.

| # Sample ID | Sex | Age (yr) | Time since HIV diagnosis (yr) | Viral Load (copies/ml) | CD4 Cell Count (cells/µl) | Time on ART-virologically suppressed (yr) | ART regimen |
|---|---|---|---|---|---|---|---|

Not available in the preprint version.

### Extended Data Figure 1

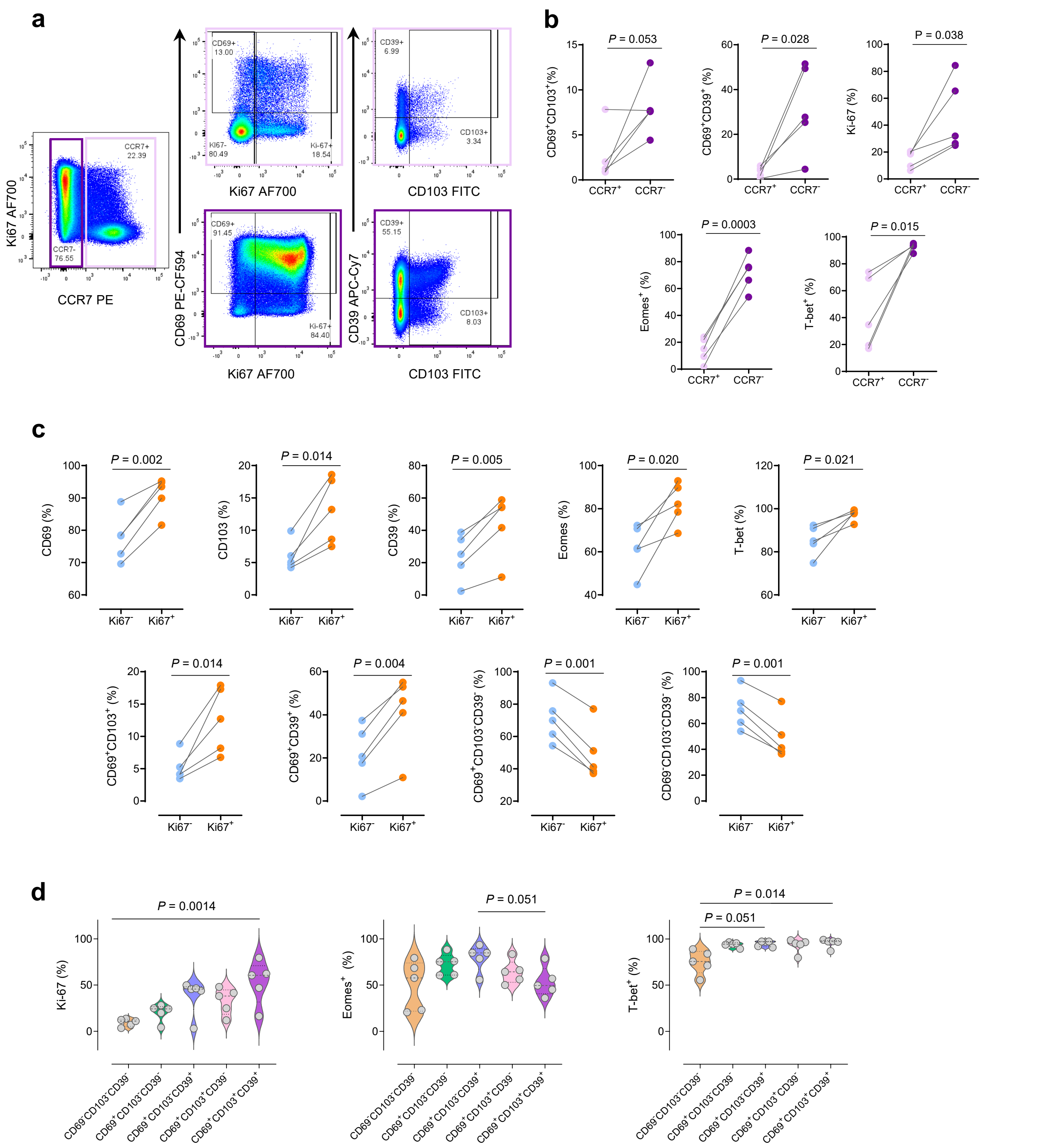


**Extended Data Fig. 1. Induction of CD69⁺ CD103⁺/CD39⁺ Ki-67⁺phenotypes in effector CCR7⁻CD8⁺T cells after sequential IL-15/TGF-β1 treatment of PBMC from ART-suppressed patients. a,** Representative flow cytometry plots after stimulating PBMC from a PWH (#M36, **Extended Data Table 1**) with sequential IL-15/TGF-β1 treatment comparing the expression levels of CD39, CD69 and CD103 between the $CCR7^+$ (top) and $CCR7^-$ (bottom) $CD8^+$T cell fractions. **b-c**, Paired comparisons of the frequencies of subsets and individual proteins (indicated in the y axis) within (**b**) $CCR7^+CD8^+$T cells (dark green dots) compared to $CCR7^-CD8^+$T cells (light green dots) or (**c**) Ki-$67^-$ (blue dots) versus Ki-$67^+$ (purple dots) in PWH (n=5, #M36-40-41 and PB01-02, **Extended Data Table 1**). **d**, Expression of Ki-67, Eomes and T-bet in $CCR7^-CD8^+$ T cells based on the indicated subsets of interest with combined expression of CD69 and/or CD103 and/or CD39 in the same PWH as in (**b-c**). Statistical comparisons for (**b**) and (**c**) were performed by a parametric two-tailed paired T test, while for (**d**) were performed using Friedman test with post hoc Dunn's correction for multiple comparisons.

### Extended Data Figure 2

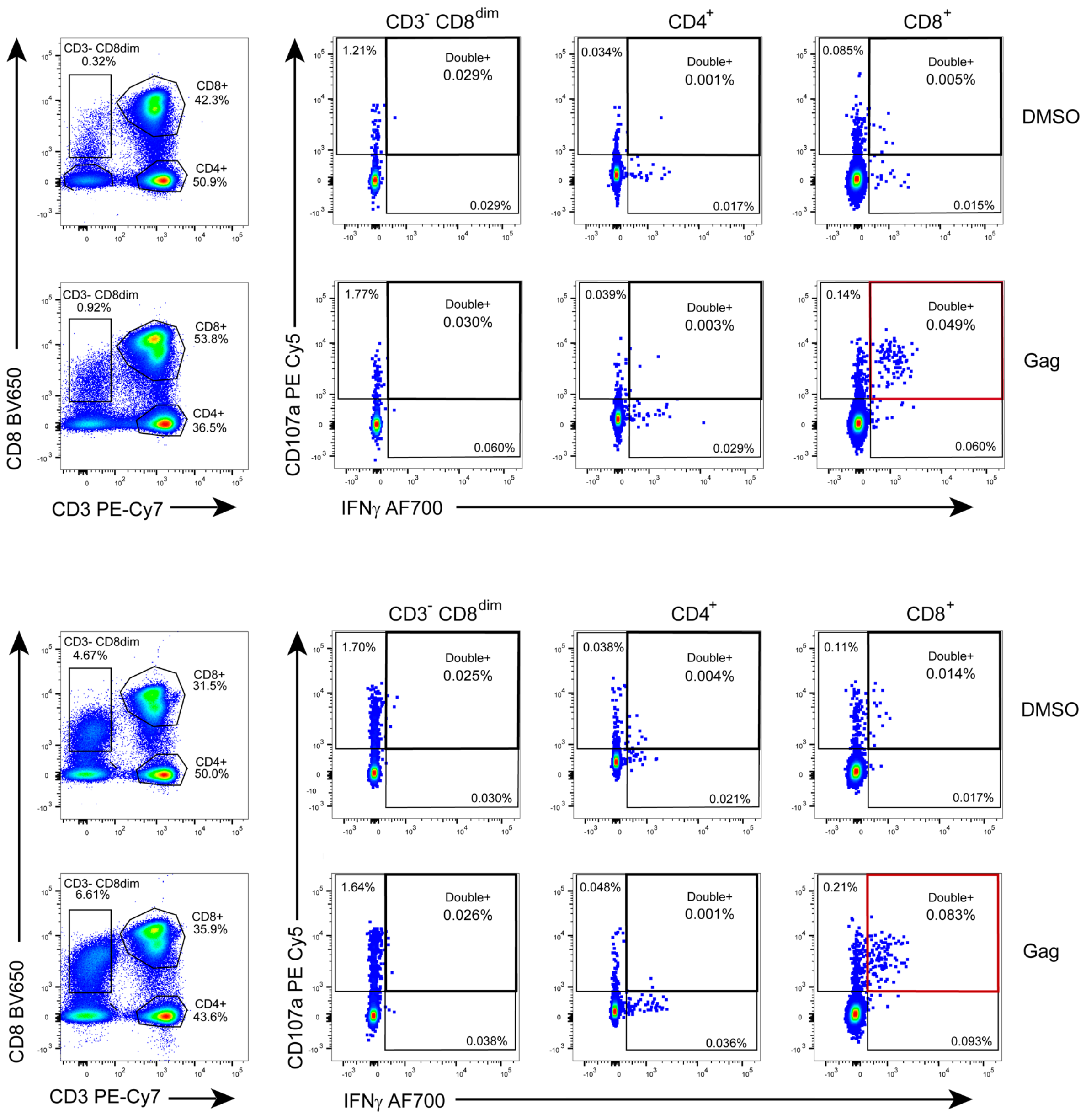


**Extended Data Fig. 2. Polyfunctional Gag-specific responses after sequential IL-15/TGF-β1 stimulation are restricted to the $CD8^{+}$T cell fraction.** Representative flow cytometry plots of different CD3/CD8 subsets in two PWH (#M36 top and #M39 bottom, **Extended Data Table 1**). For each sample, the expression of CD107a and IFNγ in DMSO (top row) or Gag-stimulated (bottom row) conditions is shown for $CD3^{-}$ $CD8^{dim}$, $CD3^{+}CD4^{+}$T cells and $CD3^{+}CD8^{+}$T cells (from right to left). Red square highlights double $IFN\gamma^{+}CD107a^{+}$ Gag-specific $CD8^{+}$T cells.

### Extended Data Figure 3

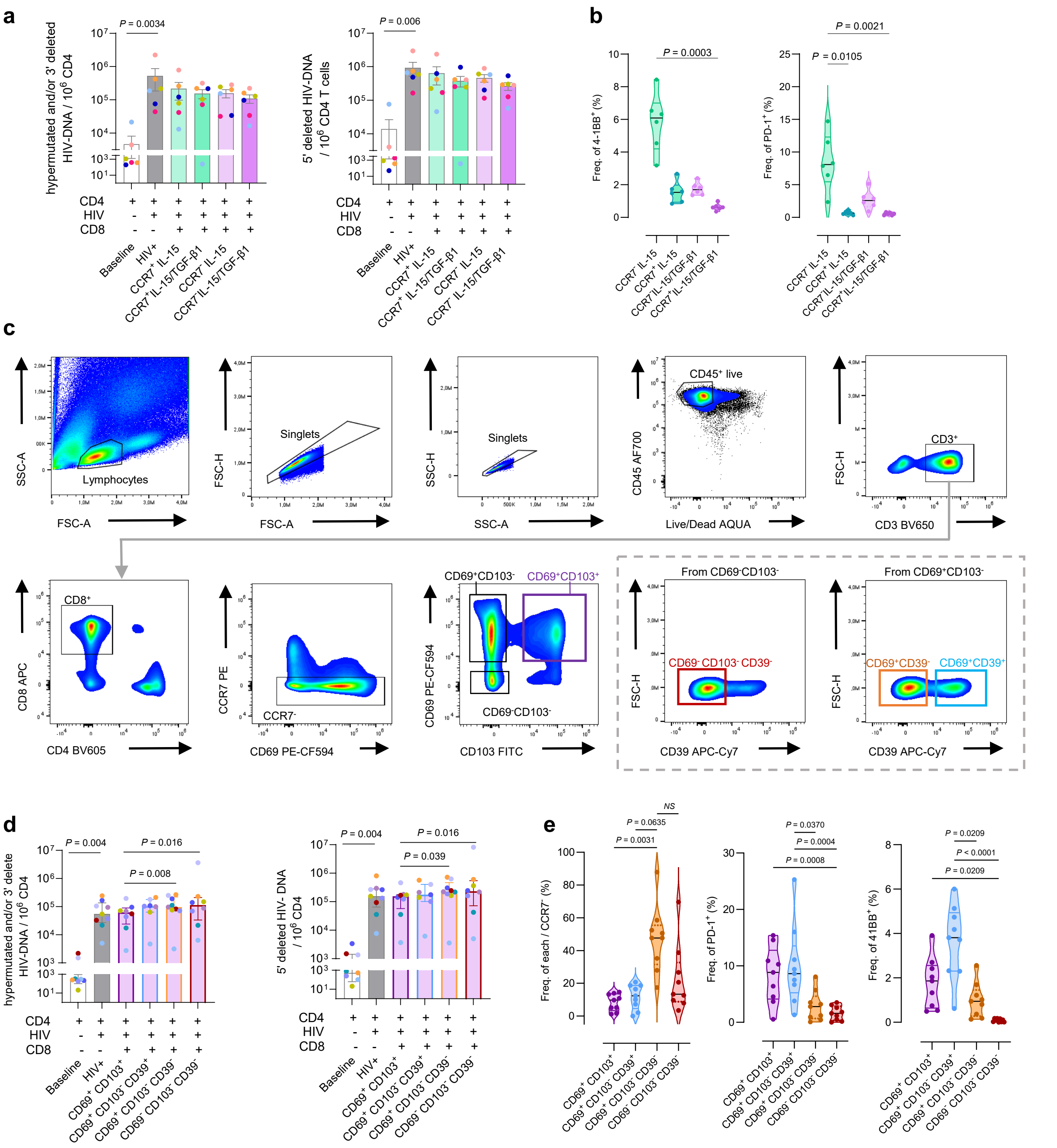

**Extended Data Fig. 3. Phenotype of sorted CD8⁺T cell subsets used for functional analyses and elimination of defective HIV forms. a,** Bar chart showing molecules of defective forms of HIV-DNA per million of CD4⁺T cells after *ex vivo* re-infection of samples from PWH co-cultured in the presence of indicated autologous CD8⁺T cell subsets from stimulated IL-15 (in green) or IL-15/TGF-β1 (in purple) PBMC (n=6, #PB02, PB21-25; **Extended Data Table 1**). **b**, Violin plots showing the frequencies of PD-1⁺ or 41BB⁺ cells for each CCR7⁻ and CCR7⁺ fraction used in (**a**). **c**, Gating strategy used to purify four CCR7⁻CD8⁺T cell subsets from IL-15/TGF-β1 stimulated PBMC, namely CD69⁺CD103⁺ (in purple), CD69⁺CD103⁻CD39⁺ (in blue), CD69⁺CD103⁻CD39⁻ (in orange) and CD69⁻CD103⁻CD39⁻ (in red), tested for their capacity to eliminate intact proviruses in co-cultures (*see* **Figure 2i**) and defective forms, as shown here in (**d**). **d**, Bar chart showing molecules of defective forms of HIV-DNA per million of CD4⁺T cells from PWH after *ex vivo* re-infection and co-culture with the indicated CD8⁺T cell subsets from IL-15/TGF-β1 stimulated PBMC sorted as shown in (**c**) (n=9, #M36, M40, PB02-03, PB07, PB12-PB15, **Extended Data Table 1**). **e**, Violin plots showing the frequencies of PD-1⁺ or 4-1BB⁺ cells for each subset tested in (**d and Figure 2i**). For (**a**) and (**d**) each colour represents an individual sample, and statistics were performed using two-sided nonparametric Wilcoxon matched-pairs signed rank test for group comparison or, for (**b**) and (**e**) the non-parametric paired Friedman test with Dunn's multiple comparisons. Bars or lines and error bars in violin plots represent median and interquartile ranges.

### Extended Data Figure 4

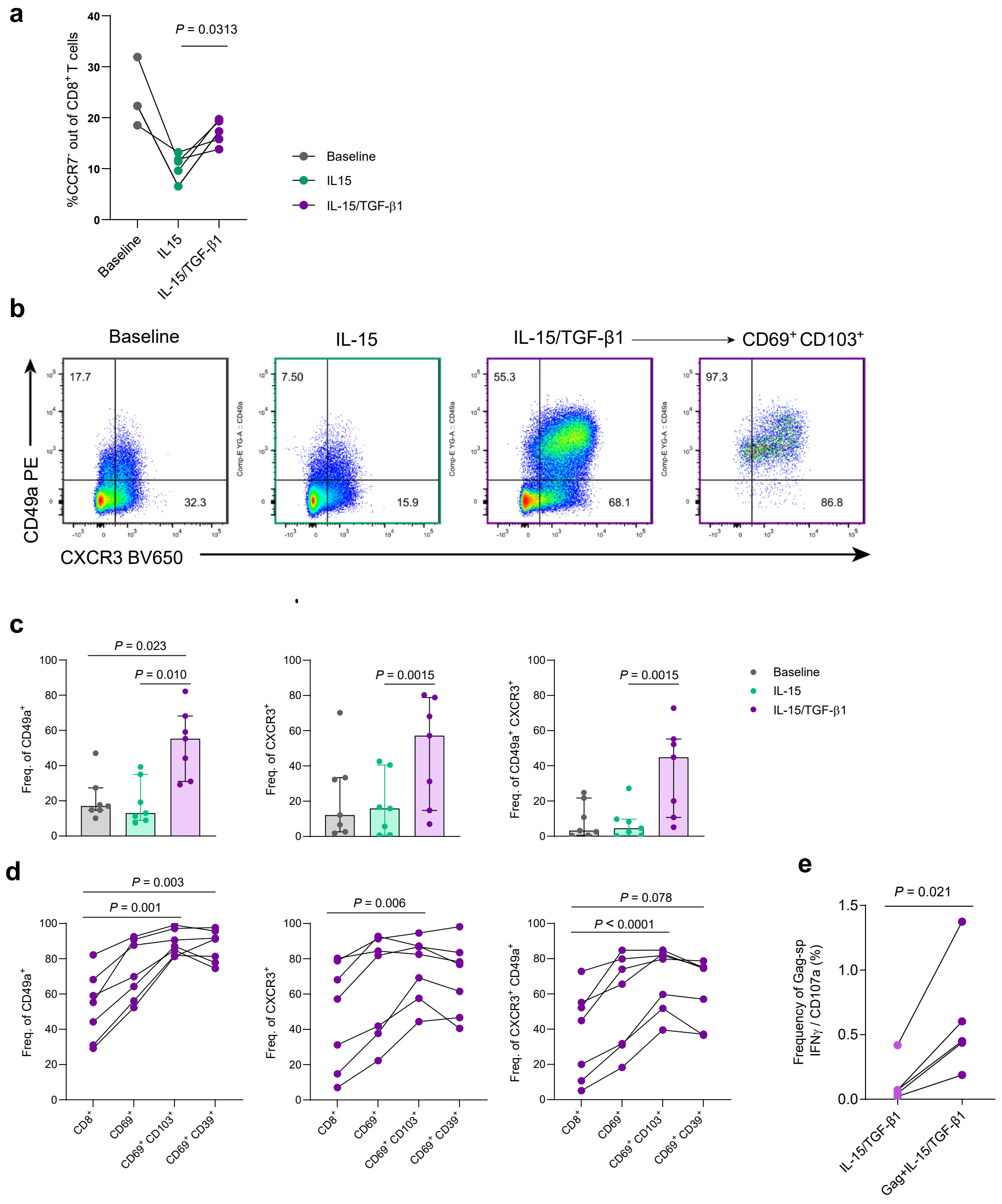


**Extended Data Fig. 4. Frequency of CCR7⁻ and of CXCR3, CD49a CD8⁺T cells after sequential treatment. a,** Paired comparison of the frequency of CCR7⁻ out of CD8⁺T cells present at baseline, IL-15 or after sequential treatment of PBMC from ART-suppressed patients (PWH, n=5, #M36, M47-49, PB16-17; Extended Data Table 1). **b,** Representative flow cytometry plots of CD49a and CXCR3 expression in CD8⁺T cells from a patient left untreated, treated with IL-15 or with IL-15/TGF-β1, from which the CD69⁺CD103⁺ fraction induced by the sequential treatment is also shown. **c,** Bar charts comparing the frequency of CD49a⁺, CXCR3⁺ or CD49a⁺CXCR3⁺ in CD8⁺T cells from each condition, as indicated in the legend (n=7). **d,** Paired comparisons of the frequency of CD49a⁺, CXCR3⁺ or CD49a⁺CXCR3⁺ in total CD8⁺T cells, CD69⁺, CD69⁺CD103⁺ and CD69⁺CD39⁺ CD8⁺ T cells from the IL-15/TGF-β1-treated condition of PBMC from the same patients as in (**c**). **e**, Paired comparison of the net frequency (DMSO-subtracted) of double IFNγ/CD107a Gag-specific CD8⁺T cells in PBMC from PWH (n=5) stimulated with sequential IL-15/TGF-β1 treatment (light purple dots) or with the inclusion of Gag during IL-15 stimulation of the sequential treatment (dark purple dots). Statistical analyses were performed using two-sided non-parametric Wilcoxon matched pairs signed rank test for (**a** and **e**), two-sided non-parametric Friedman with Dunn's multiple comparison test for (**c**) and parametric two-tailed paired T test for (**d**). PBMC-samples: baseline (grey), IL-15+Gag (green) and sequential IL-15+Gag/TGF-β1-stimulation (purple).

### Extended Data Figure 5

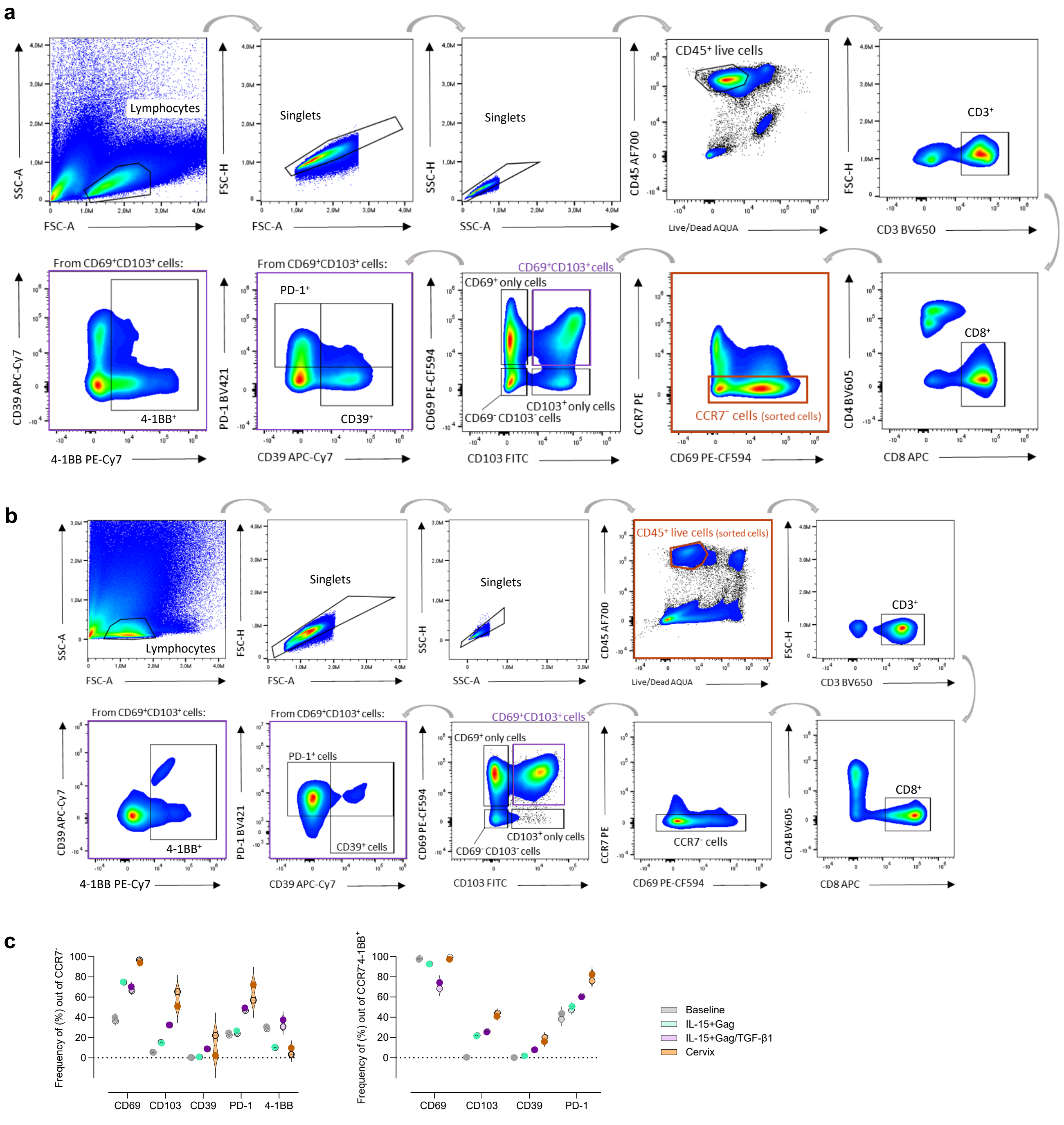


**Extended Data Fig. 5. Gating strategy and phenotype of blood and cervical samples sequenced. a** and **b,** Gating strategy used for the isolation of CD8⁺CCR7⁻ T cells from PBMC (**a**) or the isolation of live CD45⁺ lymphocytes from cervical tissue (**b**) of samples derived from patient #M47 (**Extended Data Table 1**) in which single cell RNA sequencing analyses were performed. **c,** Expression of indicated molecules within total CD8⁺CCR7⁻T cells or within CD8⁺CCR7⁻41BB⁺T cells of the PBMC conditions and cervix. PBMC-samples: baseline (grey), IL-15+Gag (green) and sequential IL-15+Gag/TGF-β1-stimulation (purple) after overnight DMSO (empty circles) or Gag (full circles) stimulation. Cervical samples (orange): control cervix (DMSO) and #M47-cervical sample (Gag).

### Extended Data Figure 6

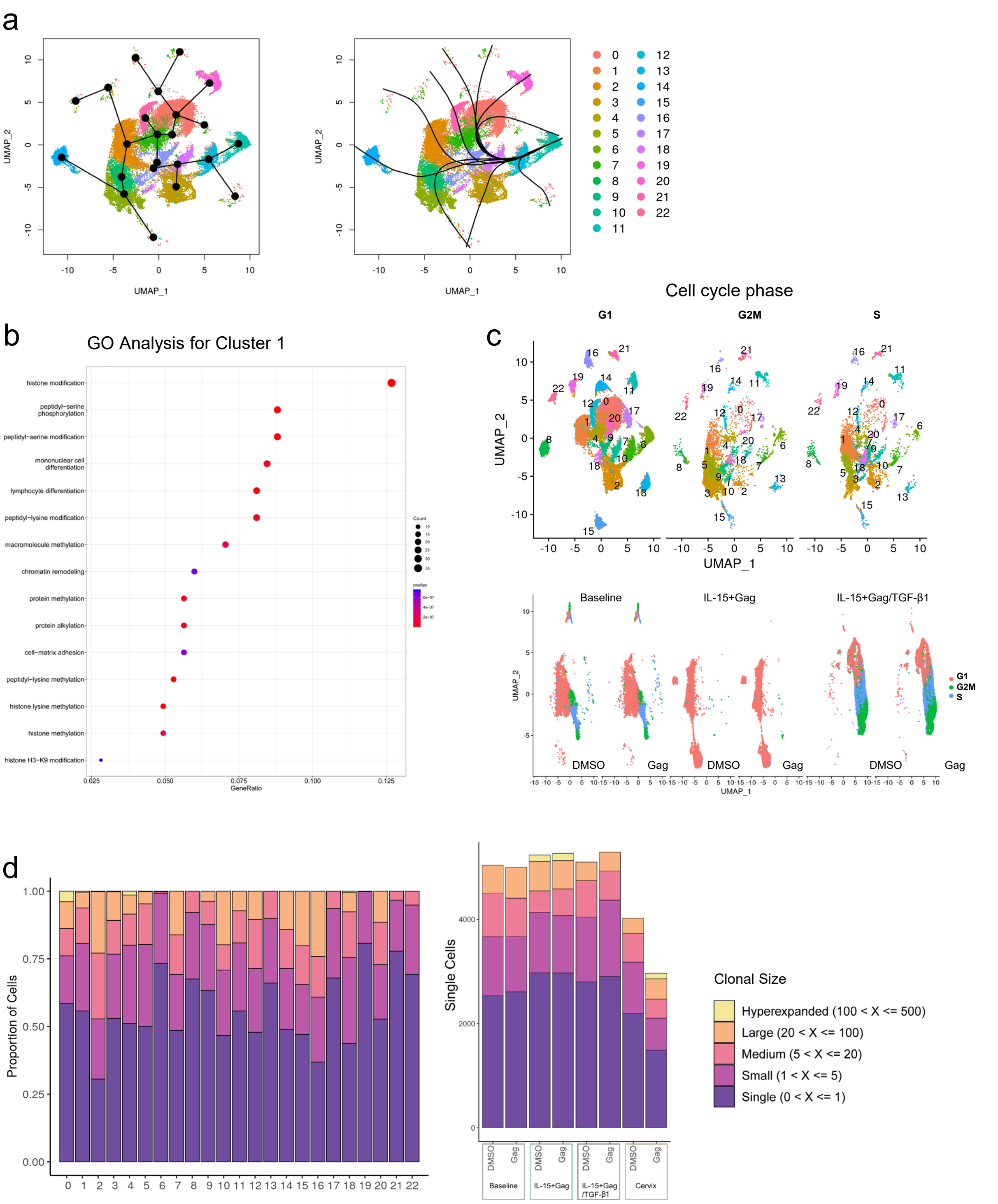


**Extended Data Fig. 6. Trajectory, gene ontology, cell cycle phase and clone size after sequential treatment. a,** Trajectories visualized on top of UMAP representations with cells colored by cluster considering six PBMC conditions (#M47). **b**, Plot showing the summary of the Gene Ontology (GO) analysis obtained for cluster 1 considering six PBMC conditions (#M47). **c,** UMAP plots showing the segregation of clusters by cell cycle (top) and cell cycle by sample (bottom), considering the six PBMC conditions (#M47). **d**, Plots showing relative frequencies (left) and cell numbers (right) of occupied repertoire space calculated for each cluster considering the six PBMC conditions (left) or for each sample considering all eight samples (#M47, six from blood and two from cervical tissue), categorized in different clonal sizes as indicated in the corresponding legend.

### Extended Data Figure 7

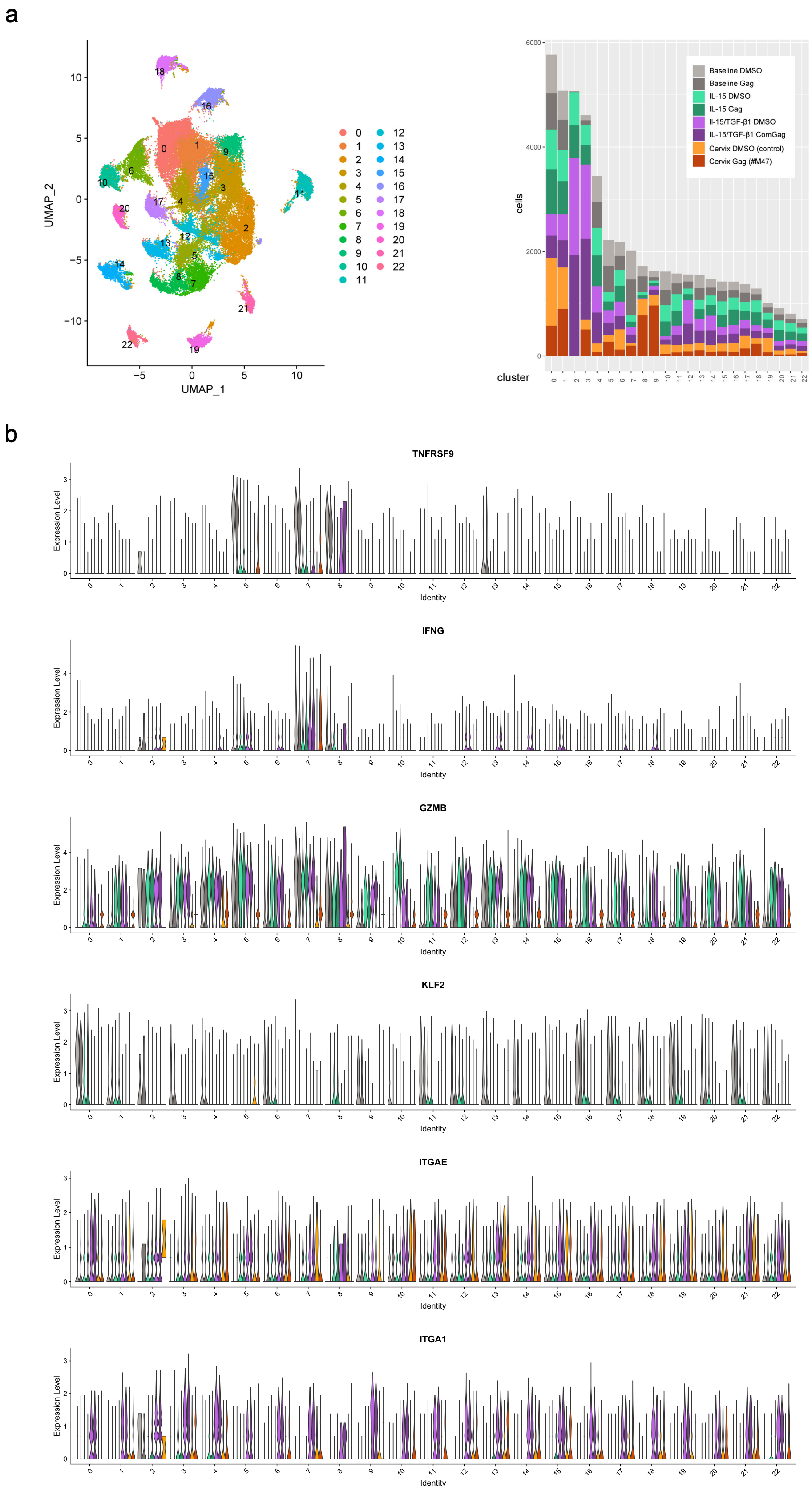


**Extended Data Fig. 7.** Cluster segregation and expression of selected genes in blood and cervical samples. a, UMAP plot showing the segregation of CD8+T cell clusters at a 0.5 resolution considering all eight samples (left) and corresponding stacked bar chart showing cell number per sample for each cluster (right). b, Violin plots showing the expression level of highlighted genes by cluster for each sample: cluster 1 to 22, considering 8 samples as shown in (a).

### Extended Data Figure 8

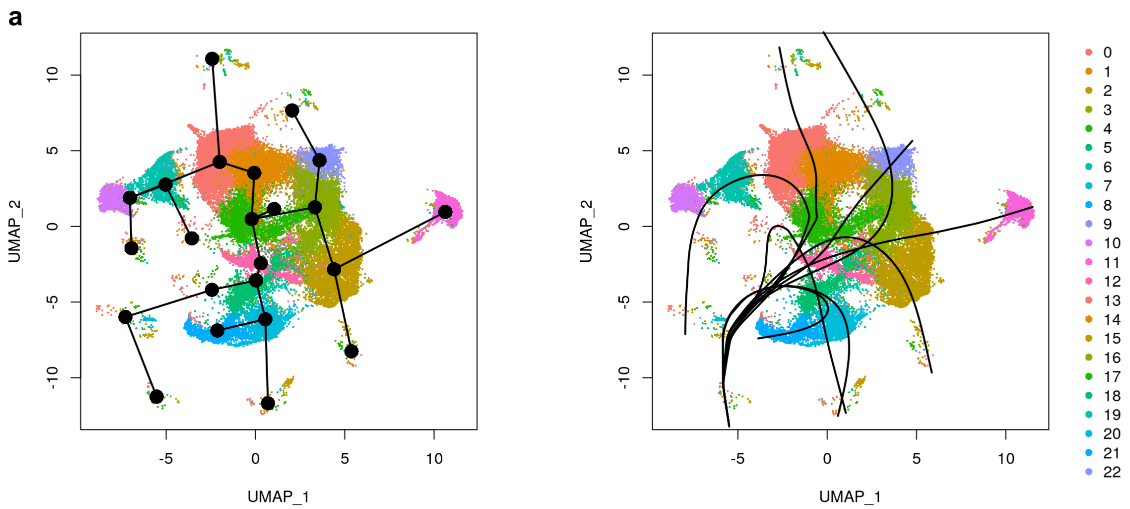


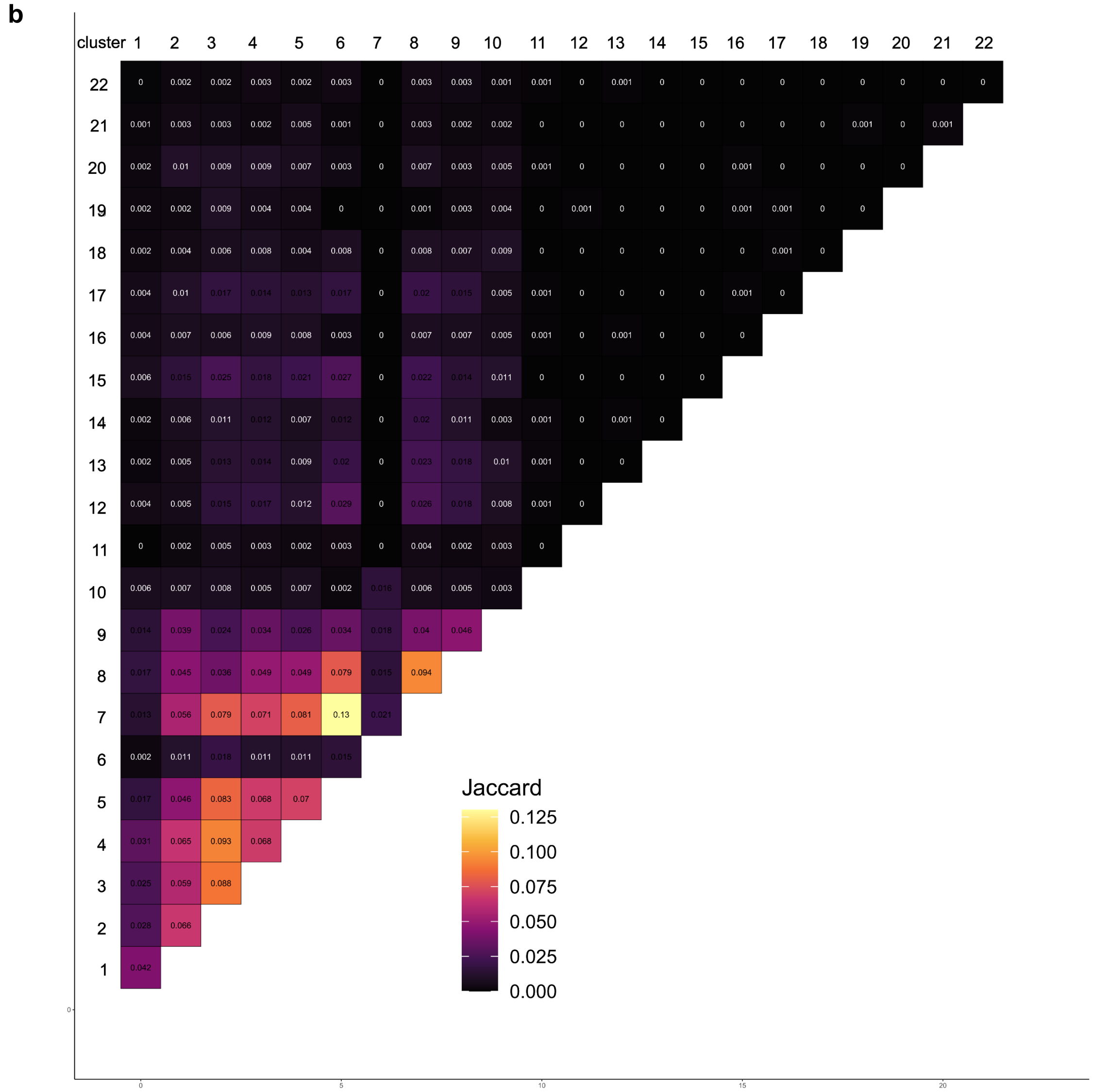


**Extended Data Fig. 8. Cluster trajectories and clonal overlap between blood and cervical samples. a,** Trajectories visualized on top of UMAP representations with cells colored by cluster considering all eight blood and cervical samples. **b**, clonal overlap was measured using Jaccard index and visualized using heatmaps.

### Extended Data Figure 9

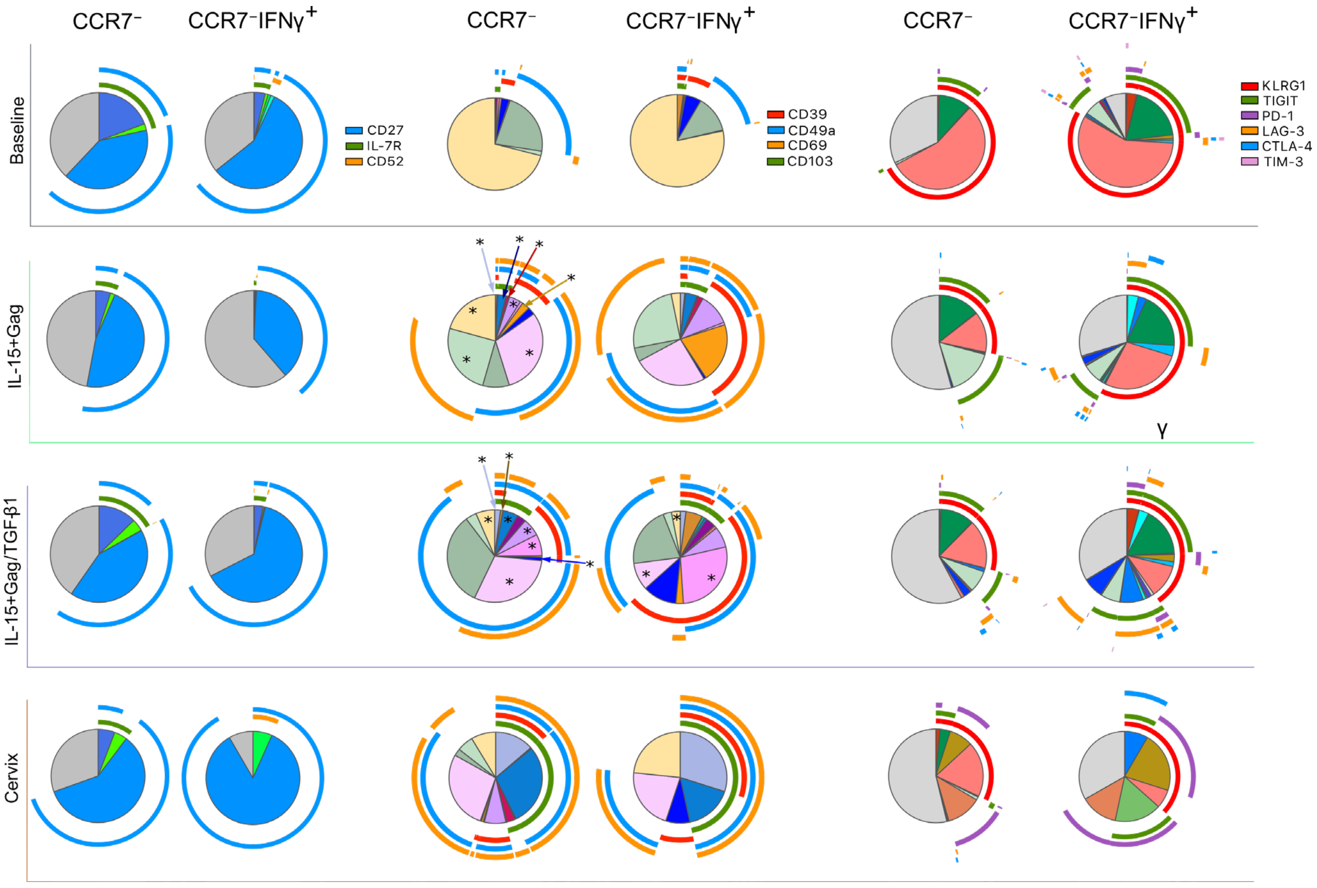


**Extended Data Fig. 9. Phenotype of effector CD8⁺T subpopulations induced by sequential cytokine treatment compared to blood and cervical samples. a,** Pie charts displaying the proportion of CCR7⁻ (left) and CCR7⁻IFNγ⁺ (right) CD8⁺T cell expressing multiple proteins analysed by boolean gates in study samples from PWH (n=8, #M36, M47-M51, PB16-17; **Extended Data Table 1**): PBMC: Baseline, IL-15+Gag and IL-15+Gag/TGF-β1, and in four matched cervical samples (bottom, #M48-51). Each colourful fraction inside each pie chart indicates a proportion of cells expressing a given phenotype indicated by the outside arches (as indicated in the corresponding legend). Statistical significance measured by one-way ANOVA with Dunn's correction are indicated with an asterisk for a given fraction (*$P < 0.05$).

### Extended Data Figure 10

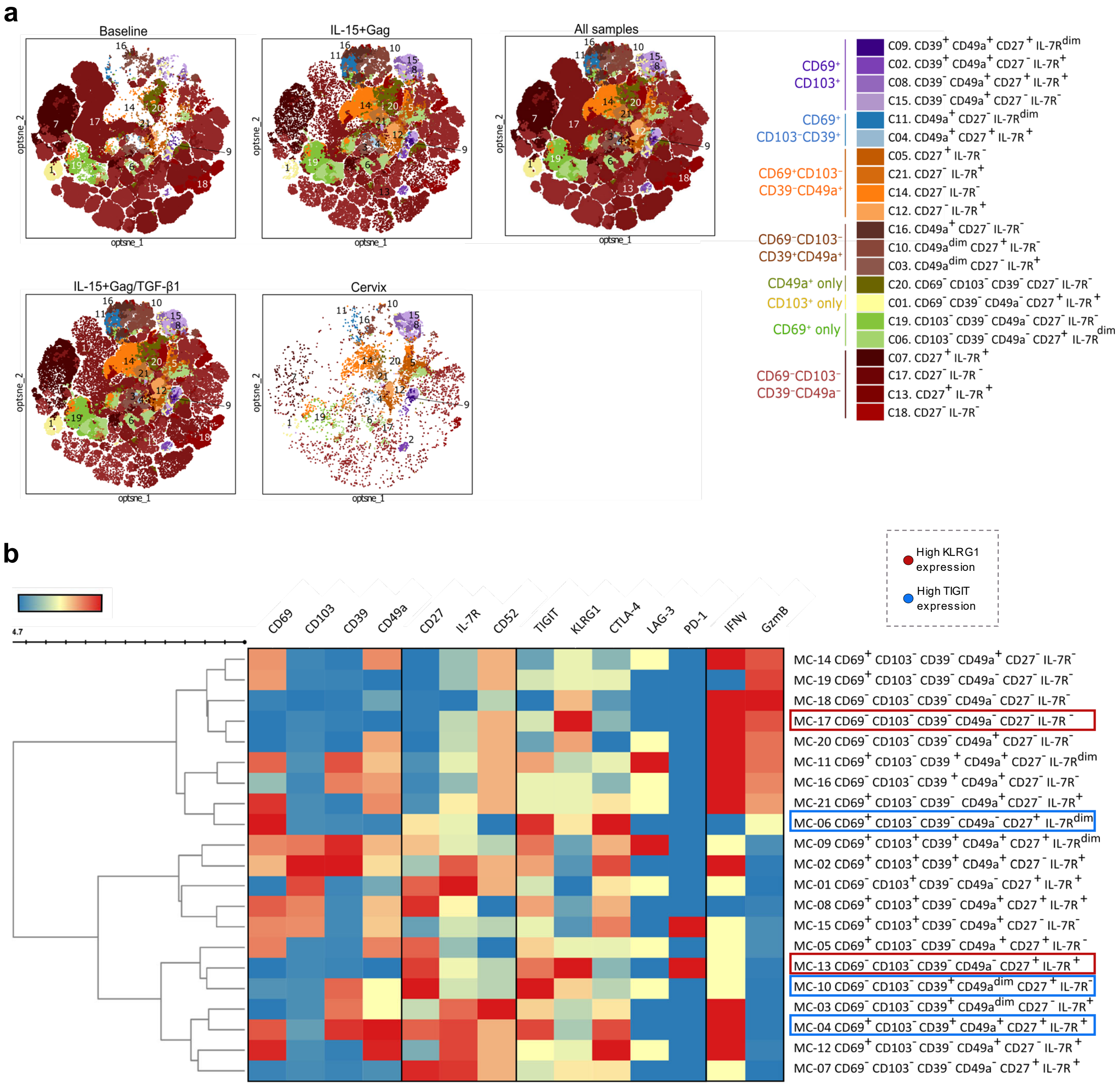


**Extended Data Fig. 10. Phenotype of effector CD8⁺T subpopulations induced by sequential cytokine treatment compared to blood and cervical samples. a,** Opt-SNE plot displaying the distribution of the 21 cell clusters identified based on the expression of CD69, CD103, CD49a, CD39, IL-7R and CD27 within CCR7⁻CD8⁺ T cells from all PBMC conditions and four matched cervical samples from the same patients as in (**a**). Legend indicates the phenotyping markers for which each cluster is enriched. **b**, Heatmap illustrating Mean Fluorescence Intensity (MFI) of all the molecules analysed across the 21 clusters identified.

**Extended Data Table 2.** Overview of the samples and immune cell composition tested in the humanized mouse model.

| # Patient ID | Mouse Code | Condition | PBMC cytokine stimulation | Cells administrated (Millions $-10^6$) | | | Ratio CD4:CD8 |
|---|---|---|---|---|---|---|---|
| | | | | $CD4^+$T cells | Non-T non-NK fraction | $CD8^+$T cells | |
| PB02 | A* | Control | None | 5 | 9 | 0 | 1.6:1 |
| | **B** | **Control** | **None** | **5** | **9** | **0** | |
| | **C** | **IL-15** | **IL-15** | **5** | **9** | **3** | |
| | **D** | **IL-15/TGF-β1** | **IL-15/TGF-β1** | **5** | **9** | **3** | |
| PB15 | E† | Control | None | 7 | 3.2 | 0 | 2.2:1 |
| | **F** | **Control** | **None** | **7** | **3.2** | **0** | |
| | **G[$]** | **Control** | **None** | **7** | **3.2** | **0** | |
| | H* | Control | None | 7 | 3.2 | 0 | |
| | **I** | **IL-15** | **IL-15** | **7** | **3.2** | **3.12** | |
| | **J** | **IL-15/TGF-β1** | **IL-15/TGF-β1** | **7** | **3.2** | **3.12** | |
| M44 | **K[$]** | **Control** | **None** | **6.45** | **0** | **0** | 1.7:1 |
| | **L** | **Control** | **None** | **6.45** | **3.6** | **0** | |
| | **M[$]** | **Control** | **None** | **6.45** | **3.6** | **0** | |
| | Nº | Control | None | 6.45 | 3.6 | 0 | |
| | O* | IL-15 | IL-15 | 6.45 | 3.6 | 3.8 | |
| | **P** | **IL-15** | **IL-15** | **6.45** | **3.6** | **3.8** | |
| | **Q** | **IL-15/TGF-β1** | **IL-15/TGF-β1** | **6.45** | **3.6** | **3.8** | |
| | **R** | **IL-15/TGF-β1** | **IL-15/TGF-β1** | **6.45** | **3.6** | **3.8** | |
| | S* | IL-15/TGF-β1 | IL-15/TGF-β1 | 6.45 | 3.6 | 3.8 | |
| | T* | IL-15/TGF-β1 | IL-15/TGF-β1 | 6.45 | 3.6 | 3.8 | |
| PB18 | Aº | Control | None | 4.5 | 1.46 | 0 | 1:1 |
| | **B** | **Control** | **None** | **4.5** | **1.46** | **0** | |
| | **C** | **IL-15** | **IL-15** | **4.5** | **1.46** | **4.5** | |
| | **D** | **IL-15** | **IL-15** | **4.5** | **1.46** | **4.5** | |
| | Eº* | IL-15/TGF-β1 | IL-15/TGF-β1 | 4.5 | 1.46 | 4.5 | |
| | **F** | **IL-15/TGF-β1** | **IL-15/TGF-β1** | **4.5** | **1.46** | **4.5** | |
| PB05 | **A[$]** | **Control** | **None** | **7.63** | **1.18** | **0** | 1:1 |
| | Bº | Control | None | 7.63 | 1.18 | 0 | |
| | **C[$]** | **IL-15** | **IL-15** | **7.63** | **1.18** | **7.63** | |
| | **D+** | **IL-15** | **IL-15** | **7.63** | **1.18** | **7.63** | |
| | **E+** | **IL-15/TGF-β1** | **IL-15/TGF-β1** | **7.63** | **1.18** | **7.63** | |
| | Fº | IL-15/TGF-β1 | IL-15/TGF-β1 | 7.63 | 1.18 | 7.63 | |
| PB19 | A† | Control | None | 5 | 3.44 | 0 | 1:1 |
| | C | IL-15 | IL-15 | 5 | 3.44 | 5 | |
| | E | IL-15/TGF-β1 | IL-15/TGF-β1 | 5 | 3.44 | 5 | |

| | | | | | | | |
|---|---|---|---|---|---|---|---|
| PB03 | A | Control | None | 5 | 1.24 | 0 | 1:1 |
| | B | Control | None | 5 | 1.24 | 0 | |
| | C°* | IL-15 | IL-15 | 5 | 1.24 | 5 | |
| | E | IL-15/TGF-β1 | IL-15/TGF-β1 | 5 | 1.24 | 5 | |
| | F | IL-15/TGF-β1 | IL-15/TGF-β1 | 5 | 1.24 | 5 | |
| PB20 | **A** | **Control** | **None** | **3.84** | **1.08** | **0** | 1:1 |
| | B° | Control | None | 3.84 | 1.08 | 0 | |
| | **C** | **IL-15** | **IL-15** | **3.84** | **1.08** | **3.84** | |
| | **E** | **IL-15/TGF-β1** | **IL-15/TGF-β1** | **3.84** | **1.08** | **3.84** | |

**Bold** Mice from which splenic tissue samples were used for the analysis.
† Mice which died before the endpoint of the experiment.
**°** Mice from which spleen sample at endpoint, and subsequently blood samples, were discarded due to low $CD4^{+}$T cell counts (<1000).
***** Mice from which spleen sample at endpoint, and subsequently blood samples, were discarded due to high $CD8^{+}$T cell counts (>500) in control group, or low $CD8^{+}$T cell counts (<500) in IL-15 or IL-15/TGF-β1 groups.
$^{\$}$ Mice from which peripheral blood samples at endpoint were discarded due to low CD45 (<500) or $CD4^{+}$T cell counts (<200).
Mice derived from an individual PWH-donor missing samples in a single arm (control, IL-15 or IL-15/TGF-β1 groups) due to low cell number or mouse death, were not included in the analyses (i.e. #PB19 and #PB03-derived mice groups).

### Extended Data Figure 11

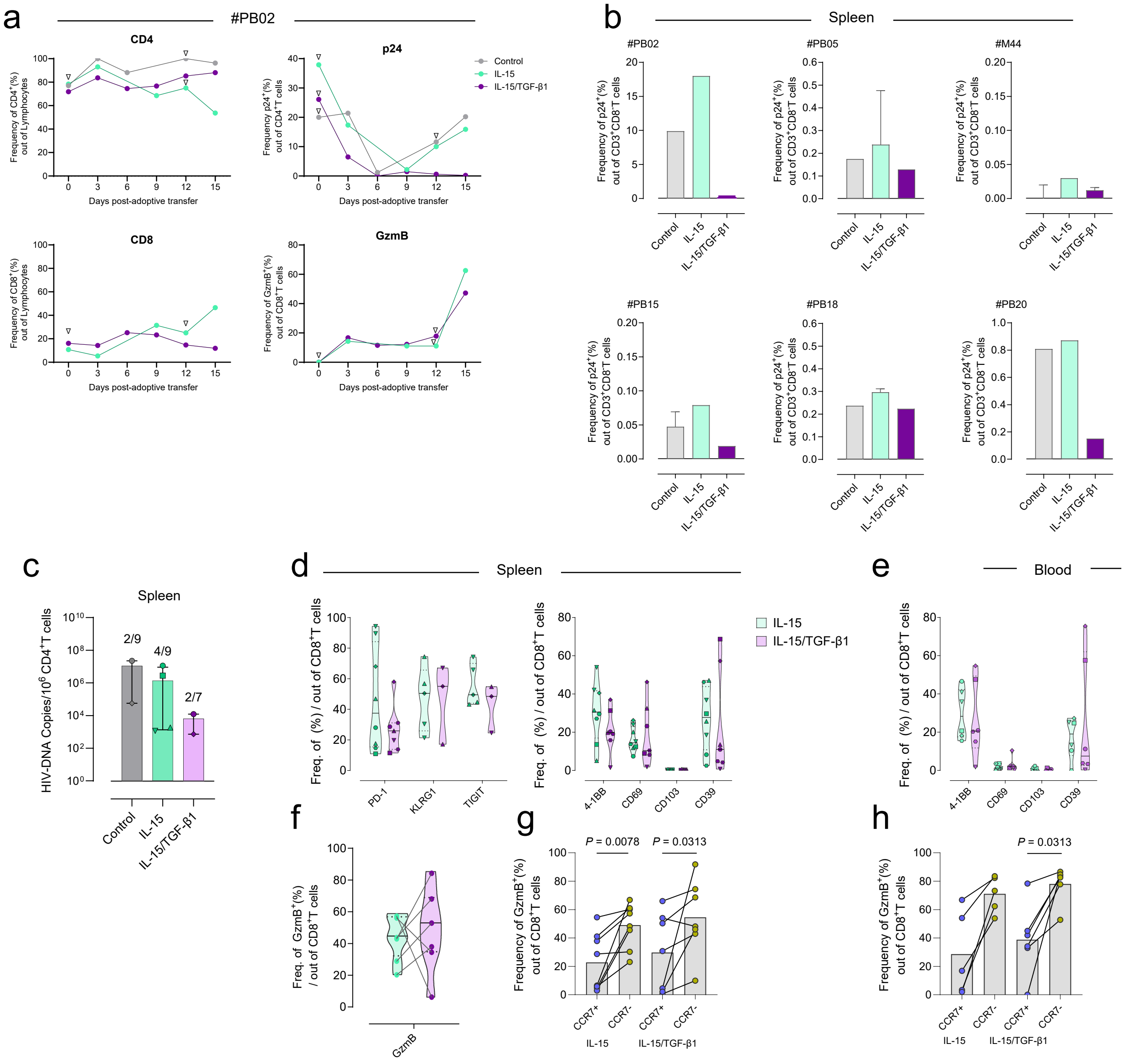


**Extended Data Fig. 10**. **Evaluation of sequentially-treated CD8⁺T cells adoptive transfer to control viral reactivation in a humanized mouse model**. **a**, Expression of CD4⁺T and CD8⁺T cells out of the total lymphocytes fraction (left), and of p24 out of CD4⁺T cells and Granzyme (Gzm) B out of CD8⁺T cells (right) from longitudinal blood samples of a single mouse per condition of the #PB02-derived group (control in grey, IL-15 in green and IL-15/TGF-β1 in purple). Samples with less than 100 CD45⁺, 50 CD3⁺ counts, or 50 CD8⁺T cells are highlighted with an inverted triangle (∇). **b**, Bar charts showing the splenic p24⁺ frequencies out of CD3⁺ CD8⁻T cells for each individual PWH-derived mouse group across the three groups of study. **c**, Bar chart depicting proviral HIV-DNA copies per million CD4⁺T cells detected in spleen for each study group. Ratios on top of each bar depict the number of samples with detectable HIV-DNA levels out of samples that could be assessed per group (corresponding to #PB02, PB05, PB15, PB18 and PB20, **Extended Data Table 1 and 2**). **d** and **e**, Frequencies of different molecules out of total CD8⁺T cells in splenic tissue (**d**) or blood (**e**) of mice from the IL-15 or the sequential treatment group. **f**, Paired comparisons of GzmB expression out of total CD8⁺ T cells in splenic tissue between IL-15 or sequentially-treated mice from the same patient-derived group performed by calculating the mean of the frequencies of the mice belonging to the same group and PWH, compared to the mean of the other PWH-group. **h** and **g**, Paired bar charts of the frequencies of GzmB⁺ cells within CCR7⁺ (blue) and CCR7⁻ (yellow-green) CD8⁺T cell fractions across IL-15 and sequential treatment mouse groups from blood (**h**) and spleen (**g**). For spleen determinations, n=8 mice for the IL-15 and n=7 for sequential treatment-group derived from 6 PWH (#M44, PB02, PB05, PB15, PB18 and PB20; **Extended Data Table 1 and 2**) except for KLRG1 and TIGIT expression with 5 samples from IL-15 and 3 from sequential mouse groups derived from 3 patients (#PB05, PB18 and PB20; **Extended Data Table 1 and 2**). Paired comparison excluded mouse samples derived from #PB05 patient due to lack of cell counts. Samples corresponding to a single PWH-derived group are indicated as follows: hexagon for #M44, circle for #PB02, triangle for #PB05, square for #PB15, inverted triangle for #PB18 and rhombus for #PB20.
